# Ventral tegmental dopamine neurons control the impulse vector during motivated behavior

**DOI:** 10.1101/2020.03.10.985879

**Authors:** Ryan N. Hughes, Konstantin I. Bakhurin, Elijah A. Petter, Glenn D.R. Watson, Namsoo Kim, Alexander D. Friedman, Henry H. Yin

## Abstract

The Ventral Tegmental Area (VTA) is a major source of dopamine, especially to the limbic brain regions. Despite decades of research, the function of VTA dopamine neurons remains controversial. Here, using a novel head-fixed behavioral system with five orthogonal force sensors, we show for the first time that distinct populations of VTA dopamine activity precisely represent the impulse vector (force exerted over time) generated by the animal. Optogenetic excitation of VTA dopamine neurons quantitatively determines impulse in the forward direction, and optogenetic inhibition produces impulse in the backward direction. At the same time, these neurons also regulate the initiation and execution of anticipatory licking. Our results indicate that VTA controls the magnitude, direction, and duration of force used to move towards or away from any motivationally relevant stimuli.

**One Sentence Summary:** VTA dopamine bidirectionally controls impulse vector and anticipatory behavior

## Introduction

The ventral tegmental area (VTA) is a midbrain region with dopamine (DA) neurons that project widely to the frontal cortex, limbic basal ganglia, and other brain regions [1]. Despite decades of research, the function of these neurons remains controversial. According to one prominent hypothesis, DA neurons encode a reward prediction error (RPE), which serves as a teaching signal that updates the value of learned associations [2–4]. Others argue that DA encodes incentive salience to motivationally relevant stimuli, or the amount of effort (or vigor) the animal exerts towards these stimuli [5–10]. Still others argue that DA neurons can multiplex different cognitive, reward, and kinematic variables [11, 12].

One potential source of ambiguity in DA function is the lack of quantitative behavioral measurements. In most studies investigating VTA DA, animals are often head-fixed, and the behavioral measures are usually limited to time stamps of licking, eye or arm movements [13–15]. Although animals in head-fixed setups often produce many movements of the head and body, these are usually ignored. In studies with freely moving animals, behavioral measures also lack spatial resolution and cannot accurately measure subtle and direction-specific movements [16]. Consequently, although DA activity has been extensively linked to the delivery of rewards and reward-predicting cues, whether these neurons are involved in the generation of movements remains unclear due to the lack of adequate measures. To improve behavioral measures in head-fixed animals, we developed a novel apparatus for measuring the head and body force generated during behavior[17]. We show that VTA DA neurons provide a precise representation of the impulse vector (force exerted over time), and regulate the direction, amplitude, and duration of force generated by the animal during motivated behavior.

## Results

In order to record the single-unit activity of VTA DA neurons, we used both *WT* mice (*n* = 3) and *DAT-Cre* mice crossed with *Ai32* mice (*n* =4), and chronically implanted them with electrode arrays (Figure 1A)[18]. *DAT-Cre* mice express Cre-recombinase only in neurons that express the dopamine active transporter (DAT), and *Ai-32* mice express Cre-dependent excitatory channelrhodopsin (ChR2). We can thus confirm that the neurons we recorded from are dopaminergic (DAT+) neurons by selectively activating VTA_DAT+_ neurons while recording their single unit activity (tagged neurons: *n* = 44, from 4 different mice: *n* = 34, 5, 4 and 1; total neurons: *n* = 298; Figures 1B-E)[19]. Despite heterogeneity in the waveform widths and firing rates [15], most optically tagged VTA_DAT+_ neurons exhibited low firing rates (FR) and wide waveform widths (mean FR = 3.99 ± 0.35 Hz; Valley Full Width at Half Maximum (FWHM) = 624.10 ± 38.72 μs; Figures 1C & 1E). Based on our optogenetic tagging results, we classified the remaining population as DA if their Valley FWHM was wider than 500 μs and their mean firing rate was less than 10 Hz (*n* = 127; Figure 1E). Using an unbiased Gaussian Mixture Model (GMM), we then functionally classified the DA neurons into three separate populations (Figures 1F-G, see below for detailed description).

**Fig. 1.**
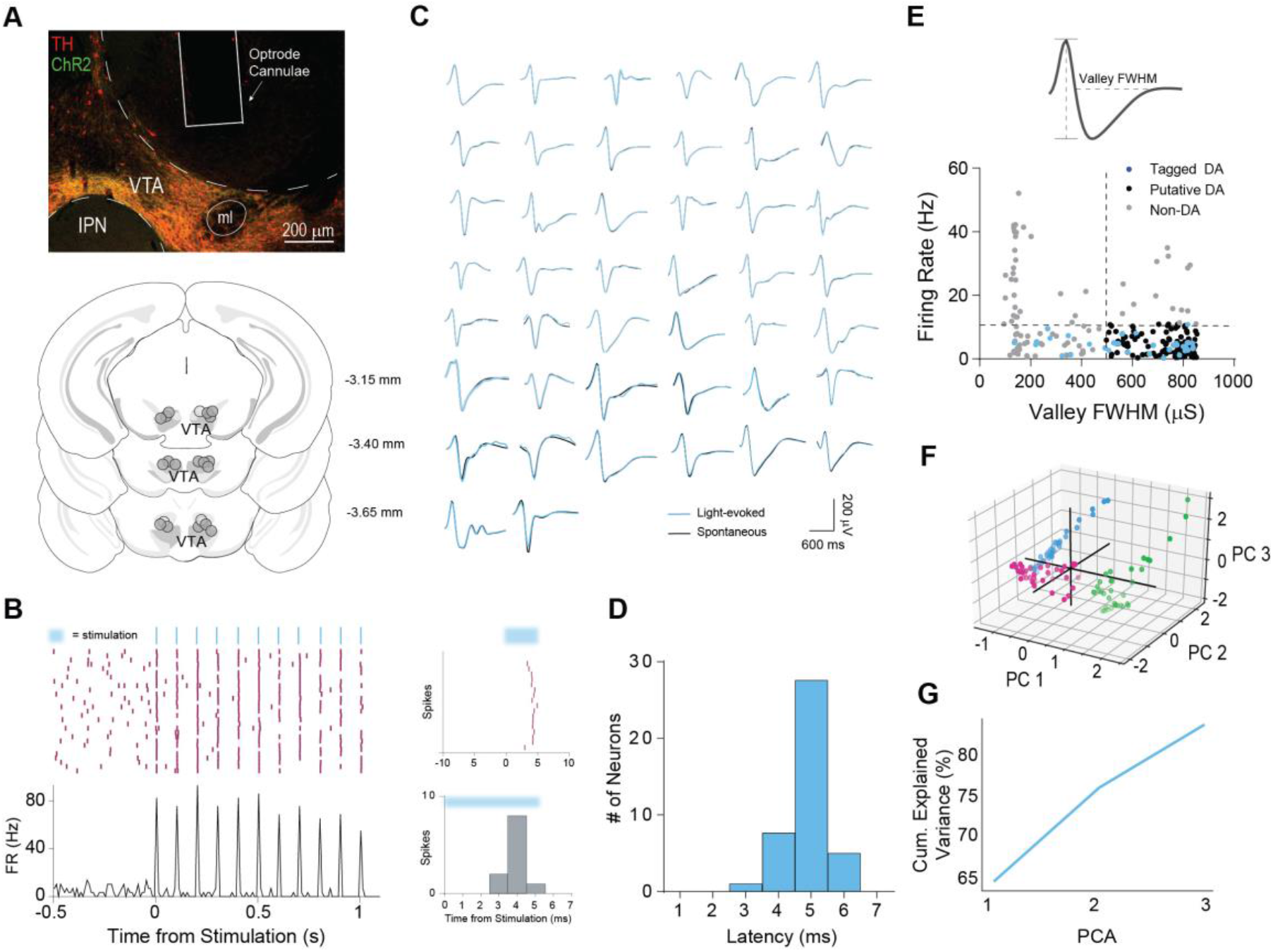
Summary of Optic Tagging and Classification of VTA DA Neurons. **A)** *Top*: Coronal section showing optrode placement along with colocalization of tyorsine hydroxylase (TH) and channelrhodopsin (ChR2) in DAT + Ai32 mice. *Bottom*: Schematic of locations of electrode implants (*n* = 7) **B)** *Left*: Representative example of a tagged VTA dopamine (DA) neuron using 10 Hz stimulation. *Right*: Zoomed in view of the same tagged neuron’s spiking showing latencies of 4-5 ms (top) and the latency of each individual spike (bottom). **C)** Fidelity analysis of tagged VTA_DAT+_ neurons showing spontaneous waveforms (black) and light evoked waveforms (blue). **D)** All tagged VTA DA neurons had a latency of ≤= 6 ms (*n* = 32). **E)** Classification for remaining DA population. The majority of tagged VTA DA neurons (*n* = 44) had a firing rate (FR) of less than 10 Hz and a valley full width at half max (FWHM) greater than 500 μs. **F)** A Gaussian Mixture Model (GMM) was applied to all VTA DA neurons based on their functional properties during the behavioral task. The graph depicts 3 clusters or populations. G) More than 85% of the variance of each population could be accounted for with 3 principal components.

To precisely quantify the forces that animals produce while restrained, we developed a head-fixation apparatus that incorporates orthogonal force-detecting load-cells coupled to the head-bar clamps (Figure 2A). We measured VTA DA activity during the performance of a fixed-time (FT) reward task, in which a drop of sucrose solution is delivered every 10 s in the absence of an explicit conditioned stimulus (Figure 2A). After 7 to 14 days of training, the mice exhibited clear anticipatory licking behavior prior to the onset of reward (Figure 2B). Close inspection of the signals produced by the force sensors revealed the wide diversity of movements generated by the head fixed mice. In addition to producing anticipatory licking and active consummatory behavior characteristic of the FT schedule [20], animals generated forces that varied continuously in their duration and amplitude in specific directions (Figures 2B & S1). The movements of the mice closely coincided with the duration of anticipatory lick bouts and exhibited stereotyped patterns during licking (Figure 2B). At the onset of the anticipatory lick bout, the mice typically exerted force in the forward direction, as well as pushed their head down and to one side to position themselves optimally in front of the reward spout (Figures 2B & S1). At the time of reward delivery, the mice at first either froze or moved backward very briefly, and then exerted a large push forward resulting in a sudden change in the force sensor signals. As registered by the force sensors, there was a general forward force exerted by the animals toward the reward spout (Figures 2B & S1). At the end of the consummatory lick bout, we usually detected a steep drop in force, reflecting a slight backward movement (Figure 2B). Not surprisingly, the sensor with the greatest force amplitude and most consistent pattern corresponded to the forwards and backwards direction (F/B sensor; Figure S1). We therefore focused on F/B movements in our analysis.

**Fig. 2.**
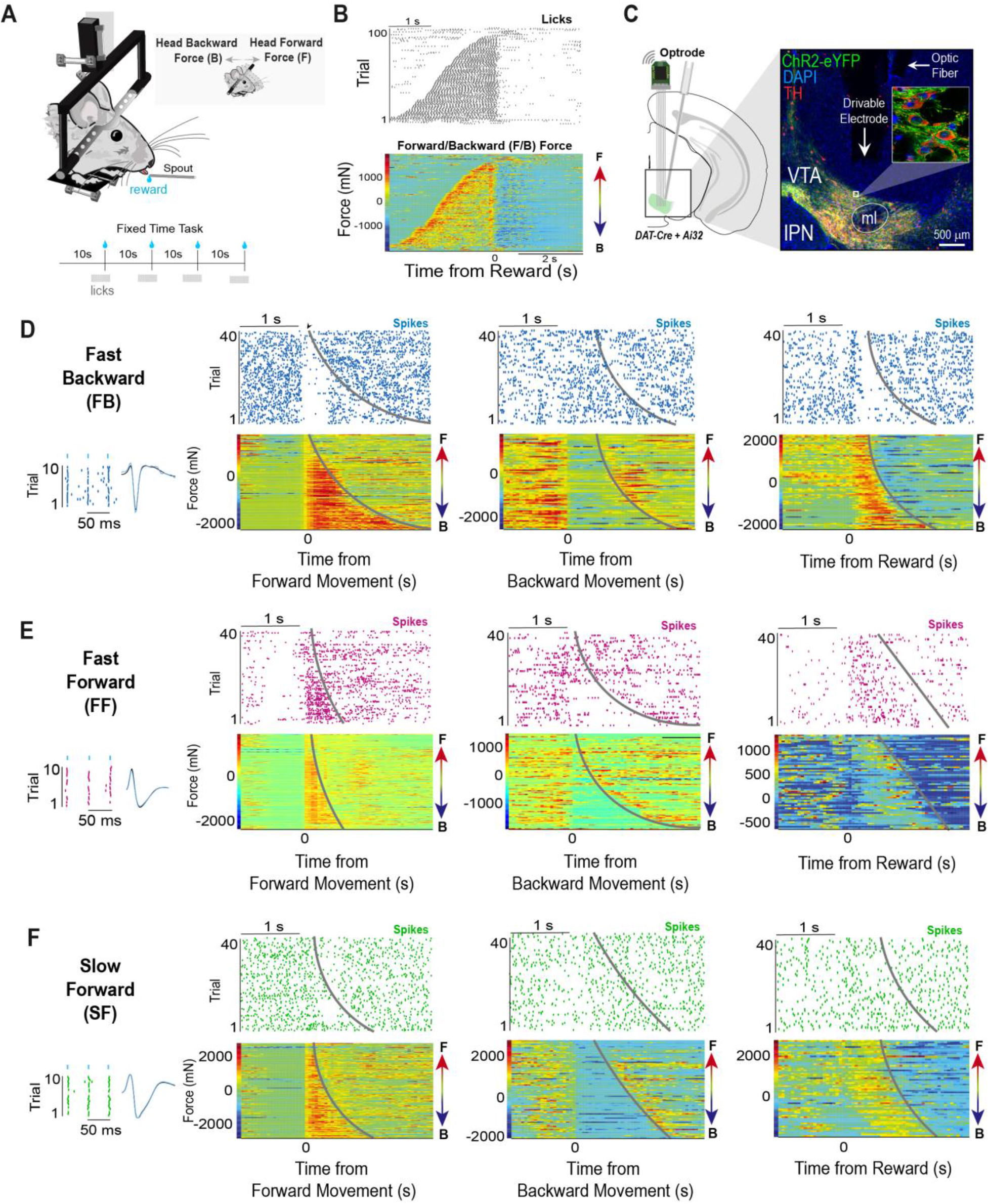
The VTA contains three types of dopamine neurons that are distinct in their responses to reward and direction of movement. **A)** *Top*: Schematic representation of novel head-fixation apparatus with five orthogonal force sensors. Inset shows the movement axis measured by the forward/backward (F/B) force sensor. *Bottom*: Mice were trained on a fixed-time reinforcement schedule, where they received a 10% sucrose reward every 10 seconds. **B)** Mice exhibited anticipatory licking before reward delivery. *Top*: Representative raster plot of licks aligned to reward. *Bottom*: F/B force sensor reading during the same behavior shown above. Trials in both panels are sorted according to the duration of the lick bout. **C)** Schematic and coronal section showing optrode placement into the VTA. Inset shows ChR2-infected neurons also contain tyrosine hydroxylase (TH). **D)** Representative optically tagged fast-backward (FB) VTA DA neuron is inhibited during forward movement, increases its firing rate during backward movement, and displays a short-phasic burst (< 200 ms) at reward delivery. **E)** Representative optically tagged VTA DA fast-forward (FF) neuron increases its firing rate when aligned to forward movement outside of reward, is inhibited during backward movement, and displays a long-phasic burst (> 200 ms) following reward that is sustained throughout the forward movement. **F)** Representative optically tagged VTA DA slow- forward (SF) neuron increases its firing rate during forward movement, but is inhibited during backward movement, and displays little or no burst at reward. Trials in all panels are sorted according to the duration of F/B force.

We found three functional populations of VTA DA (DAT+) neurons that were distinct in their relationships to movement and reward. The first population typically displayed a short phasic burst (50 - 200 ms) at the time of reward (Figure 2D). These neurons *decreased* firing for forward movement but *increased* firing for backward movement, as indicated by the force sensors (Figure 2D; Fast-Backward (FB) neurons, total neurons: *n* = 44, tagged neurons: *n* = 23). A second population of neurons typically displayed a longer phasic burst (200 ms - 1 s in duration) at the time of reward (Figure 2E). In contrast to the FB DA neurons, these neurons *increased* their firing rate during forward movement, and *decreased* their firing rate during backward movement (Figure 2E; Fast-Forward (FF) neurons, total neurons: *n* = 32, tagged neurons: *n* = 17). Finally, a third population of DA neurons exhibited firing rates that *increased* during forward movement and *decreased* during backward movement, like FF neurons.

However, they were distinct from the FF group in that they showed little firing rate modulation at the time of reward (Figure 2F). Overall, their firing rate modulation reflected slower changes of force over time (Slow-Forward or SF neurons, total neurons: *n* = 35, tagged neurons: *n* = 4). We used the combination of responses to forward movements, backward movements, and reward to classify each DA neuron that we recorded. To further verify and validate our classification, we concatenated and scaled the response profiles of all VTA DA neurons around each of these events, then performed a principal component analysis using a GMM, which found 3 distinct clusters corresponding to our 3 functional groups (Figures 1F-G, Figure 3).

**Fig. 3.**
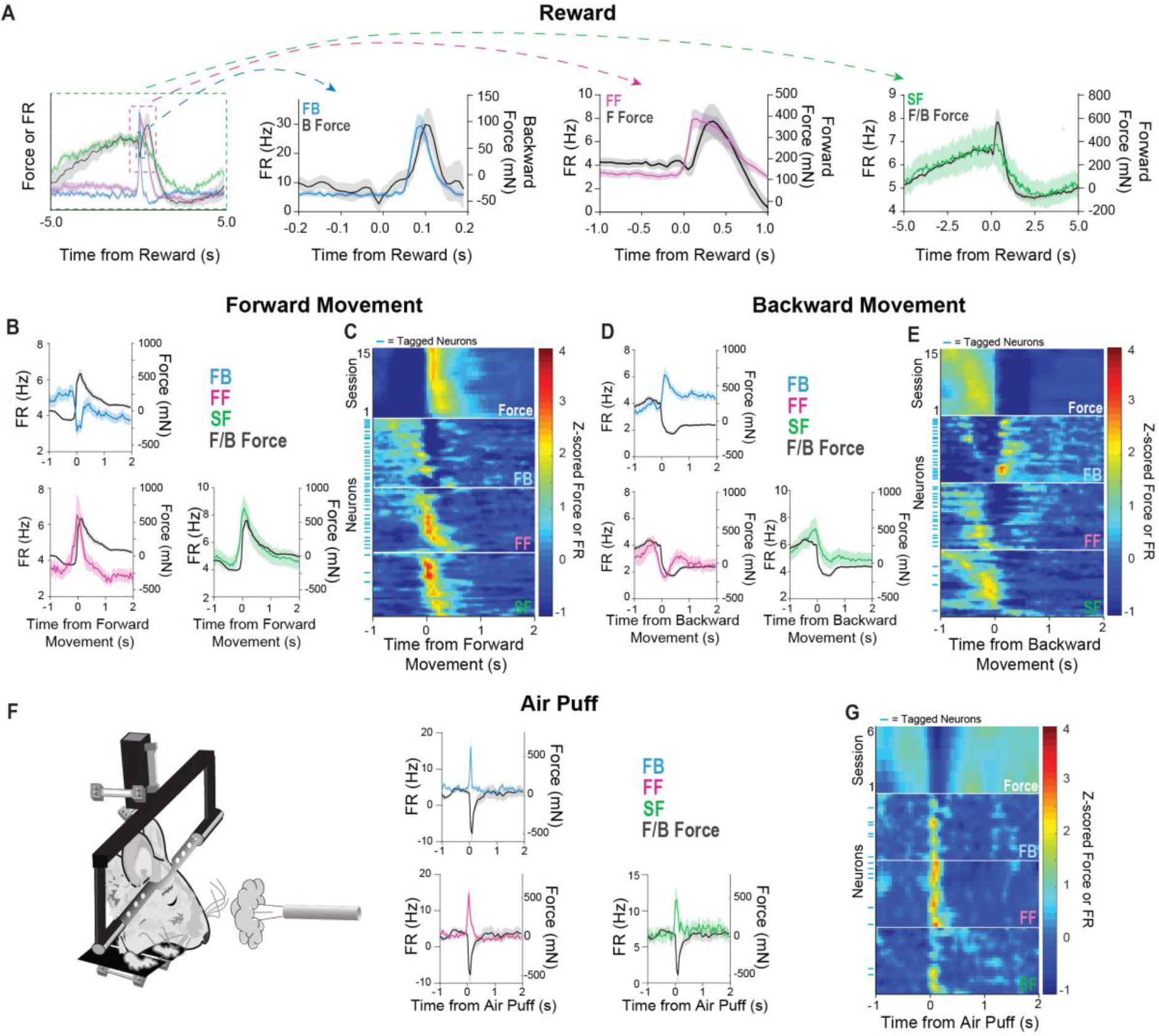
Three populations of VTA dopamine neurons can be distinguished based on direction, magnitude, and time scale of force generation. **A)** Each population of VTA DA neuron is responsive to force on different time scales and direction of movement when aligned to reward. The FB population scales with small backward forces. The FF population scales with fast forward force. The SF population scales with slower force modulation. *Left*: all populations plotted on the same time scale. *Right*: three different time scales are used on the x-axis to reveal clearly the key relationship between neural activity of each population of DA neurons and the associated force change. **B**) Average FB, FF, and SF population activity aligned to onset of forward movement outside of immediate reward delivery. The FB population decreases its firing rate during forward movement, whereas the FF population increases its firing rate. Similar to the FF population, the SF population increases its firing rate during forward movement. **C**) Heatmaps of F/B force and DA populations aligned to forward movement outside of reward times. FB neurons are inhibited during forward force generation. FF and SF neurons increase their firing rates during the duration of forward force. Blue tick marks on the left denote optically tagged neurons. **D**) Average FB, FF, and SF population activity aligned to onset of backward movements outside of immediate reward delivery. The FB population increases its firing rate during backward movement, whereas the FF population decreases its firing rate. Similar to the FF population, the SF population decreases its firing rate during backward movement. **E**) Heatmaps of F/B force and DA populations aligned to backward movement outside of reward times. FB neurons increase their firing rate for the duration of backward force. FF and SF neurons are inhibited for the duration of backward force. Blue tick marks on the left denote optically tagged neurons. **F**) *Left*: Schematic of air puff trials during the task. *Right*: Average FB, FF, and SF population activity aligned to air puff. All three populations increase their firing rate at the time of air puff. **G**) Heatmaps of F/B force, and all DA populations aligned to air puff. Blue tick marks on the left denote optically tagged neurons. Error denotes SEM.

DA neural activity usually preceded movement onset after reward (Figure S2). However, the three cell populations showed distinct relationships with force measures. After reward delivery, FB neurons showed phasic activity just before the brief backward movement (latency after reward = 64.57 ± 1.76 ms; Figures 3A & S2). In contrast, the activity of FF neurons preceded the large forward force generation following reward (Figures 3A, S2 & S3; latency after reward = 107.10 ± 8.45 ms). Neurons in the SF population, on the other hand, do not significantly modulate their firing rates at the time of reward consumption. These neurons display a much slower ramping pattern that is only apparent when a much larger time window is used (Figures 3A & S3). This suggests that the SF activity could contribute to the tonic changes seen in prior work during anticipatory approach, while the FF and FB neurons reflect the well-established phasic firing pattern of VTA DA neurons [21]. When the neural activity was aligned to forward movements outside of reward context, each population had consistent modulations that were seen around reward times. The firing rates of the FB population decreased, while the FF and SF population increased (Figures 3B-C). In contrast, the opposite patterns were observed when the neural activity was aligned to backward movement. The FB population’s firing rate increased, while the FF and SF population’s decreased (Figures 3D-E). These firing patterns during reward, forward movements, and backward movements were also seen early in training, indicating they are not responses that develop as a function of learning (Figure S3).

Although we have shown a clear relationship between force generated and DA activity, it is unclear whether these force signals are found only during reward-guided behavior. It is well established that many VTA DA neurons also respond to aversive stimuli [14, 22–24]. To examine how each DA population responded to an aversive stimulus, an air puff was delivered in some sessions to the face of the animal at random times during the task. All mice also showed very characteristic force changes: they first moved backwards, then returned to their normal position and remained motionless after the initial startle response (Figures 3F-G). Unlike during appetitive behaviors, the air puff elicited burst firing in all three populations of DA neurons (Figures 3F-G). SF and FB populations are activated first, followed by the FF population much later (Figure S2). This activation pattern reflects opponent populations producing forces in opposite directions. In other words, the aversive stimulus appears to generate conflicting force vectors, which resulted first in backward movement (due to the FB population being activated first), then forward movement followed by no movement or freezing. This interpretation is supported by our observation that the air puff produced a quick startle response that is followed by freezing behavior and no movement in any direction.

Our results show for the first time that the phasic components are responsible for fast force changes, whereas the slow component is responsible for slow force changes. Strikingly, we found that the firing rates of all three populations of DA neurons can be explained by a single variable, namely force generated over time *(impulse [Newton-seconds(J)]);* Figure 4A). We calculated the impulse resulting from each individual movement and counted the number of spikes detected in each DA neuron during that movement. This resulted in a precise linear relationship between the number of spikes of each population of VTA DA neurons and the impulse generated in a specific direction (Figure 4B). For the FB population, the number of spikes correlated highly with negative impulse, which corresponds to movement in the backward direction (Figure 4B). In contrast, the number of spikes produced by the FF and SF populations had high correlations with positive impulse, which corresponds to forward movement. These correlations were weaker when considering peak force alone (Figure S2). The FB neurons produce much less impulse over time than the FF or SF populations (Figures 4C). This is not surprising, as most natural movements are in the forward direction. In addition, the FF population appears to govern larger and faster force changes compared to the SF population, which is more responsible for generating low amplitude force changes that are typical of the force changes that occur across a longer time-window. Finally, all three types were approximately found in the same proportion (Figure 4D). Altogether, the functional contributions of each population to the generation of impulse are distinct with respect to the direction, magnitude, and the time scale of force exertion.

**Fig. 4.**
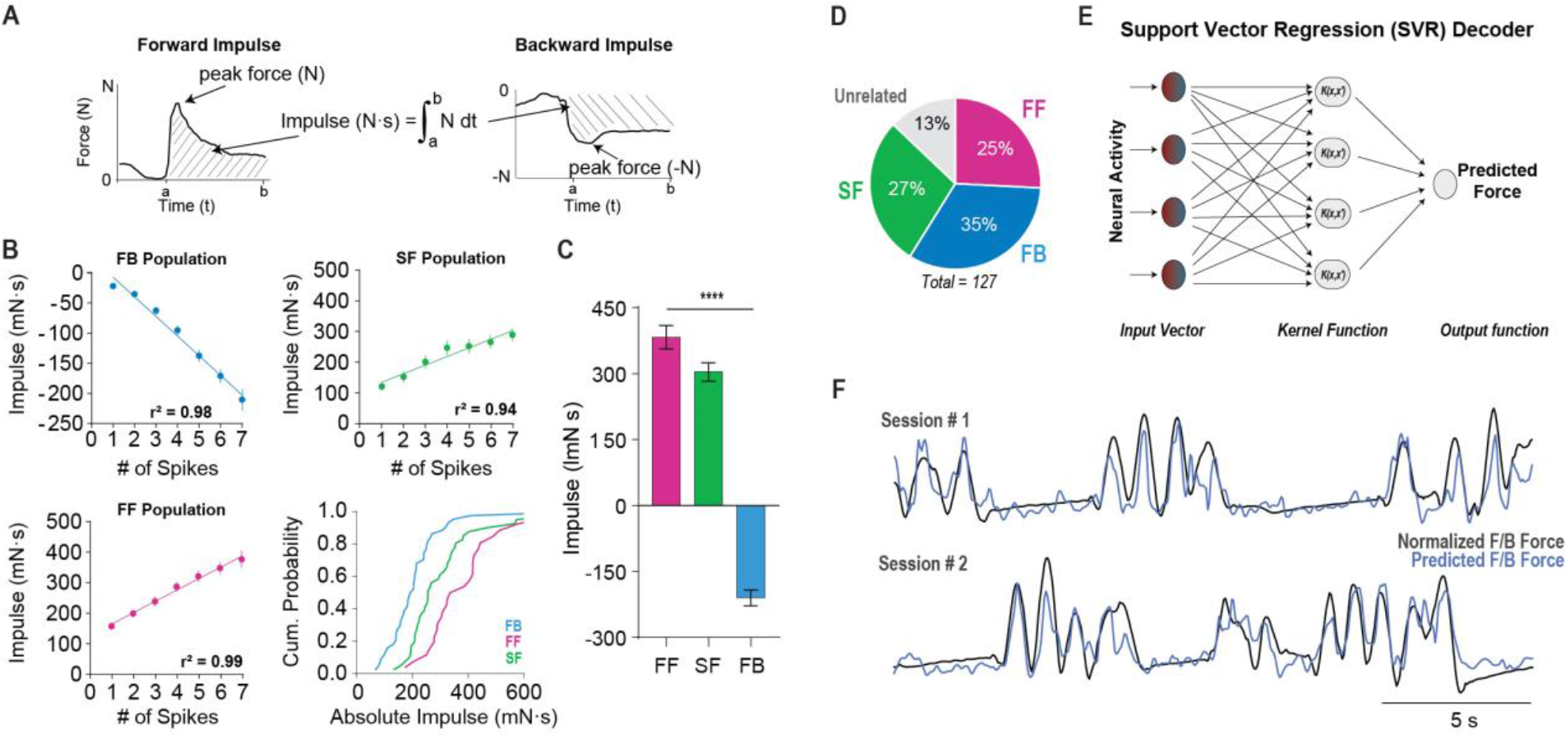
The impulse vector explains most of the variance in the activity of VTA DA neurons. **A**) Schematic of how the impulse vector is calculated. It is equal to the time integral of force (N). Negative values reflect impulse in the backward direction. **B)** High correlation with negative impulse (backwards movement) was found for the FB population, while high correlations with positive impulse (forward force) were found for the FF and SF populations. Cumulative probability shows absolute amount of impulse associated with 7 spikes (the maximum number of spikes common to all neurons during a movement). **C**) Mean impulse generated from 7 spikes from each population of neuron. There is a significant difference between all three groups (One-way ANOVA, *F*_(2,21)_ = 143.70, *p* < 0.0001. Tukey’s *post-hoc* test revealed significant differences between all three groups *p* < 0.01). **D)** Pie chart shows relative proportion of different VTA DA populations (FB: *n* = 44; FB: *n* = 33; SF: *n* = 35). **E**) Firing rates of all three populations of DA neurons can be used to predict force over time. Schematic of neural decoder used to predict forward force. **F**) F/B force was accurately decoded from all simultaneously recorded VTA DA neurons regardless of functional classification. Traces are representative 20 s examples from two different animals. Error bar indicates SEM.

Together, the activity of VTA DA neurons could be used to explain the continuous and time-varying force generated by each mouse. We used a machine learning algorithm (support vector regression) to predict the force exerted by the animal across the entire session using all simultaneously recorded DA neurons from that session regardless of their functional classification[25]. We were able to accurately decode the F/B force using only the DA population activity (Figures 4E-F & S4). Similar to the correlation analyses, the decoder was able to use the raw DA neuron population activity to determine the force generated by the animal on a moment-to-moment basis, rather than using data averaged across many trials (Figures 4E-F & S4).

Because force generation and licking bouts often co-varied, we also performed correlation analyses between VTA DA activity and licking across the entire session. All three cell populations showed significantly greater correlations with force than with lick rate (*p* < 0.0001, Figure 5A). Furthermore, when we examined all forward movements that did not coincide with a lick bout, we observed the same modulations in the activity of DA neurons (Figures 5B-C). Interestingly, when trials were sorted according to the onset of the first consummatory lick after reward, the FB neurons display several phasic bursts after reward that occurred in between licks (Figure 5D). This could be due to a strong backward force during each licking cycle, visible in the continuous force measures. To examine this possibility, we aligned the neural activity and backward force to the first and second lick that followed reward. As expected, the FB neurons increased their firing rate in a manner that closely tracked backward forces generated during consummatory licking (Figure 5E). This was also true for licking in general, not just licking at the time of reward delivery (Figure 5F).

**Fig 5.**
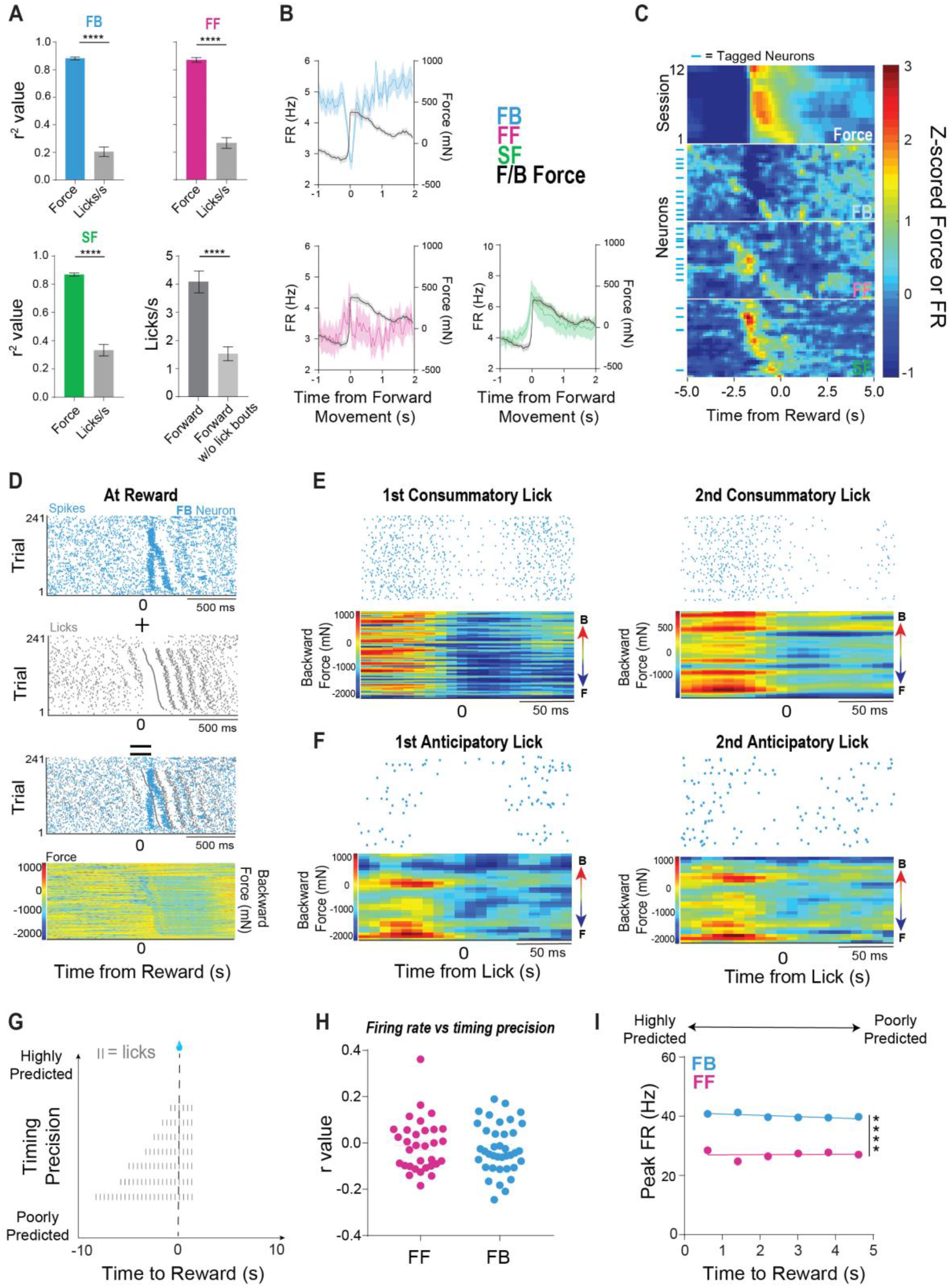
DA neurons represent force, not licking behavior. All three populations of neurons showed similar relationships with force during periods without any reward delivery or lick bouts. **A)** Correlations between neural activity and force or lick rate across the entire behavioral session. All three populations showed significantly greater correlations with force than with lick rate (*p* < 0.0001). *Bottom right*: To determine if VTA DA neural activity is still correlated with force in the absence of lick bouts, we filtered the data to examine all non-rewarded forward movements with no lick bouts. Data were filtered by removing all forward movement with lick bouts. Forward movements with lick bouts had significantly greater licks per second than forward movements without lick bouts *(p* < 0.0001). **B)** Average FF, FB, SF population activity and FB force aligned to onset of forward movement outside of reward and lick bouts. **C)** Peri-event heat map of FB force, and all neuronal populations aligned to forward movement outside of reward and lick bouts. FB neurons are inhibited for the duration of forward force. FF and SF neurons increase their firing rates during the generation of forward force. **D)** Representative optically tagged FB neuron aligned to reward and sorted according to the onset of the lick bout. When licks are overlaid with neural activity, an antiphase relationship is seen between licks and neural activity. **E)** This relationship is due to the backward force generated in between licks, which can be seen when aligned to both the first and second consummatory licks. F) This relationship is also seen when no reward is present during anticipatory licking. Error bars indicate SEM. **G)** Schematic of timing precision. The onset of the anticipatory lick bout can indicate how well the reward is predicted. Lick bouts initiated just before reward delivery reflect more precise timing and are more highly predicted. **H)** There was no correlation between the firing rate of phasic bursts for the FF and FB populations (FF: mean *r* = −0.01; FB: mean *r* = −0.02). I) The FB and FF populations have significantly different peak firing rates (Two-way ANOVA, *p* < 0.0001). However, the firing rates in each population are not modulated by reward prediction (Linear regression, FF: r_2_ = 0.0005, *p* = 0.87; FB: *r*_2_ = 0.00001, *p* = 0.10). Error bars reflect SEM.

In our task, rewards are always fully predicted [20]. Just like Pavlovian conditioning, reward delivery on a fixed-time schedule can be predicted, as indicated by anticipatory licking[26]. It is possible that phasic activity of the FB and FF populations following reward could be modulated by variability in timing, reflecting uncertainty about reward delivery time (Figure 5G). We would expect phasic firing rate to be higher when the reward is less well predicted corresponding to trials where timing is less precise. To test this possibility, we examined the firing rate of each individual reward trial and the corresponding timing of the lick bout onset before reward. We did not find a significant correlation between timing precision and post-reward DA firing rate (Figures 5H-I). There was, however, a significant difference between the peak firing rate of the FF and FB populations regardless of their timing precision, indicating they have distinct functional roles independent of a unitary prediction error signal (Figure 5I).

Because DA activity generally preceded the change in force exerted (Figure S2), it could have a causal role in force generation. To test this hypothesis, we injected Cre-dependent channelrhodopsin (DIO-ChR2) into the VTA of *DAT-Ires-Cre* (DAT::ChR2_VTA_ *n* = 5) or of *Wild-type* (WT::ChR2_VTA_ *n* = 5) mice for controls (Figures 6A & S5). We optogenetically excited (40 pulses at 20 Hz) DA neurons starting at 6 seconds before reward delivery, when animals were least likely to show anticipatory behavior. Stimulation resulted in forward force generation (Latency: 122.50 ± 7.53 ms; Figures 6B-C & S5, Movie S1). Because optogenetic stimulation cannot be limited to specific functional populations of neurons, this manipulation is expected to reflect the net effect of activating all three populations. As we found a greater proportion of neurons that increased their firing rate during forward movement (FF & SF populations; Figure 4B), a net increase in forward force generation was expected.

**Fig. 6.**
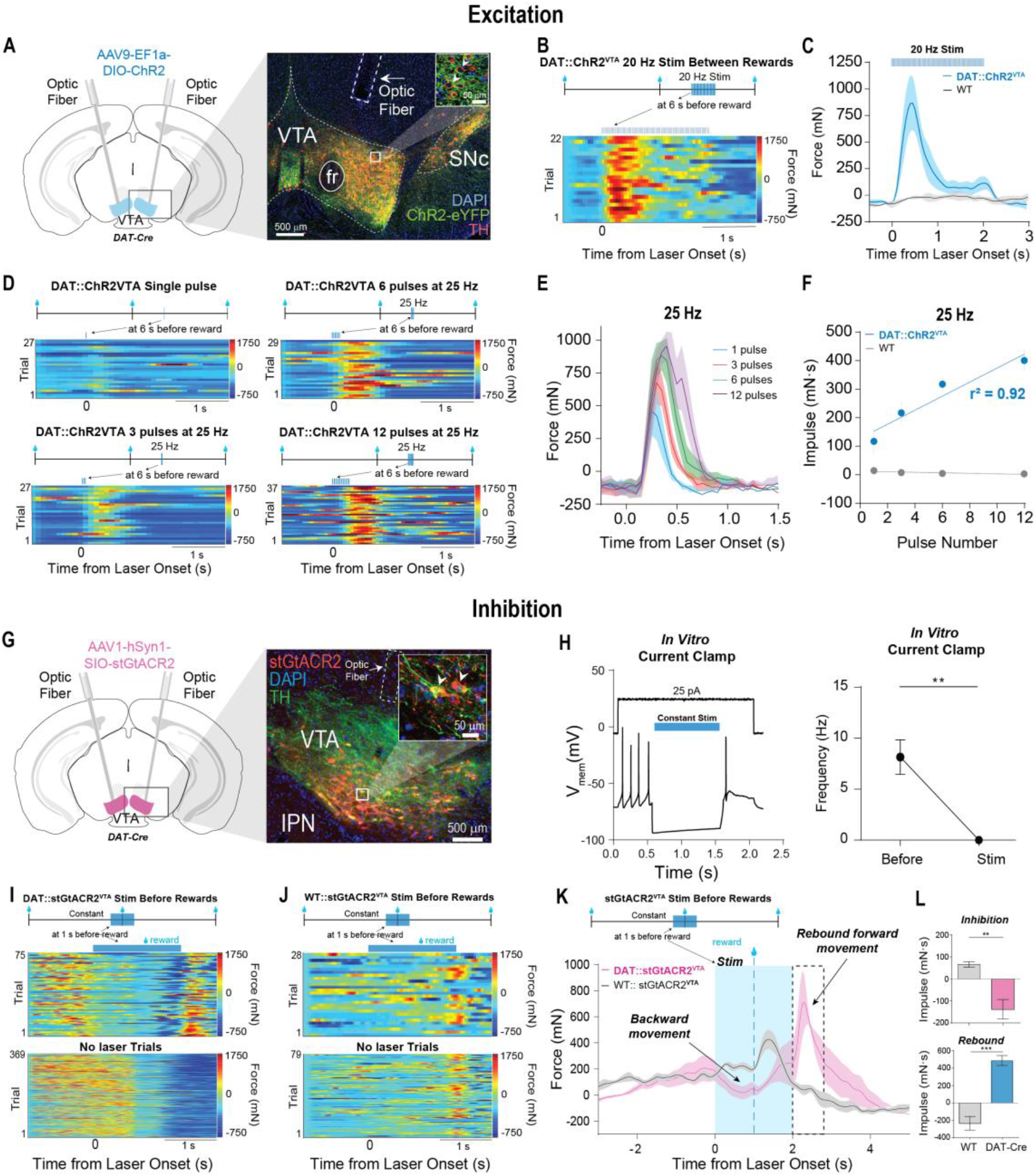
Optogenetic excitation of VTA DA neurons is sufficient to generate forward force and optogenetic inhibition of VTA DA neurons is sufficient to generate backward force, followed by rebound forward force. **A)** *Left*: Schematic illustration showing the implantation of optical fibers above VTA. *Right*: Histological confirmation of selective ChR2-expressing neurons in VTA DA neurons. Inset shows ChR2-infected neurons are TH+. **B)** *Top*: Diagram of DAT::ChR2_VTA_ excitation at 6 seconds before reward delivery. *Bottom*: Representative heatmap showing reliable forward force generation resulting from optical stimulation of VTA DA neurons at 20 Hz (5 ms pulse width) for 2 seconds. **C)** Average F/B force signals resulting from stimulation of VTA DA neurons at 20 Hz for DAT::ChR2_VTA_ (*n* = 4) and WT::ChR2_VTA_ (*n* = 5) mice. **D)** Representative heatmaps showing reliable forward force generation from 25 Hz excitation with 1,3,6 and 12 pulses (5 ms pulse duration) of light at 6 seconds before reward delivery. E) Each pulse of light produced longer duration movements and higher peak force. Plot shows forward force signals across all subjects as a function of the number of light pulses delivered at 25 Hz. F) There is a strong linear relationship between mean impulse (area under the force-time curve) and number of pulses delivered at 25 Hz in DAT::ChR2_VTA_ (*n* = 4, *p* < 0.05), but not in WT control mice (*n* = 5). G) *Left*: Schematic illustration showing injection of SIO-stGtACR2 into the VTA along with the implantation of optical fibers above VTA. *Right*: Histological confirmation of selective expression of stGtACR2 in VTA DA neurons. Inset shows colocalization of stGtACR2-infected neurons with TH-positive neurons. H) *In vitro* whole cell patch clamp recordings in current clamp mode demonstrate robust inhibition of DA neurons with stGtACR2. *Left*: Example neuron showing inhibition of VTA DA neurons using constant stimulation with 473 nm light. *Right*: Constant stimulation significantly reduced firing frequency (*n* = 5, *p* < 0.01). I) *Top*: Diagram showing DAT::stGtACR2_VTA_ inhibition (constant for 2 seconds) occurring at 1 second prior to the delivery of rewards. *Middle*: Representative heatmap showing backwards movement due to DA inhibition. Note the presence of large rebound excitation after the termination of stimulation. *Bottom*: Representative heatmap showing forward force generation prior to reward delivery on control trials without laser delivery. **J)** *Top*: Diagram showing WT::stGtACR2_VTA_ inhibition (constant for 2 seconds) occurring at 1 second prior to the delivery of rewards. *Middle*: Representative heatmap for stimulation trials for WT controls. *Bottom*: Example heatmap of control trials without laser. **K)** Average force during stimulation trials across all DAT::stGtACR2_VTA_ and WT::stGtaCR2_VTA_ animals. **L)** *Top*: There was a significant negative impulse generated by stimulation in DAT::stGtACR2_VTA_ (*n* = 4) compared to WT (*n* = 5) controls (unpaired t-test, *p* = 0.0118) during the first second of inhibition. *Bottom*: There was a significant increase in positive impulse for DAT::stGtACR2_VTA_ (*n* = 4) compared to WT (*n* = 5) controls (paired t-test, *p* = 0.0024) due to rebound excitation after inhibition ended (** *p* < 0.01; *** *p* < 0.005). Error bar indicates SEM.

It has been proposed that VTA DA neurons operate in two modes: a low firing tonic mode and a high-firing phasic mode, and that these signals can represent distinct variables at different time scales [4, 6, 12, 21, 27]. Stimulation at 20 Hz for 2 seconds could be increasing tonic DA levels as well as mimicking phasic activity. To assess the role of brief phasic firing in
force generation, we also stimulated dopamine neurons with short pulse trains (< 500ms) to mimic phasic firing activity (Figures 6D-F). Stimulation with brief trains of light resulted in consistent forward force generation (Figures 6D-F & S5). Even 1 5ms pulse of light would reliably generate movement (Figures 6D-F & S5, Movie S2). Remarkably, we also observed a strong linear relationship between the number of pulses in the stimulation and the impulse generated by the mice (Figures 6E).

Because optogenetically mimicking RPE signals in VTA DA neurons failed to produce movement in previous work [28], we tested whether forward movement generation would occur in naïve mice not previously trained on the task. Optogenetic excitation (20 Hz for 2 seconds) did not produce movement in naïve mice (Figure S5). This suggests that the motivational context is necessary for VTA DA to produce movement generation.

We next tested the effects of optogenetic inhibition on VTA DA neurons. We injected Cre-dependent soma-targeted *Guillardia theta* anion-conducting channelrhodopsin (stGtACR2), which has been shown to effectively shut down neural activity with high temporal precision[29–31], into the VTA of *DAT-Ires-Cre (*DAT::StGtACR2_VTA_, *n* =5) or of *Wild-type* (WT::StGtACR2_VTA_, *n* =5) mice for controls (Figure 6G). We also confirmed potent optogenetic silencing of DAT::StGtACR2_VTA_ neurons using *in vitro* whole cell patch-clamp recording in acutely prepared brain slices (Figures 6H & S6), demonstrating for the first time robust inhibition of DA neurons with this newly developed opsin.

We then silenced DA neurons immediately before reward delivery, when the mice normally showed high forward force production (Figure 6I). DA inhibition produced backward movement (Figures 6I & 6K-L). After stimulation ended, the mice also exhibited a rebound forward movement, most likely due to rebound excitation of VTA DA neurons from sustained inhibition, which was also seen in our *in vitro* experiments (Figures 6H, 6K-L, & S6).

Interestingly, we found that optogenetic excitation of VTA neurons 6 seconds before reward delivery (when baseline licking was at its lowest) also reliably elicited anticipatory licking in addition to exertions of forward force (Figures 7A-B, Movie S1). In contrast, optogenetic inhibition caused only a slight reduction in licking, which likely reflected a ‘floor effect’ (Figures 7E-F). Both optogenetic excitation and inhibition significantly *disrupted* anticipatory licking if stimulation occurred just before reward delivery, when the rate of licking was high (Figures 7C-D & 7G-H). However, neither optogenetic excitation nor inhibition had any effect on consummatory licking just after reward delivery (Figure S7). Together, these optogenetic stimulation results show for the first time that VTA DA neurons play a causal role in the generation of anticipatory—but not consummatory—licking, in accord with previous work [5, 20, 32].

**Fig. 7.**
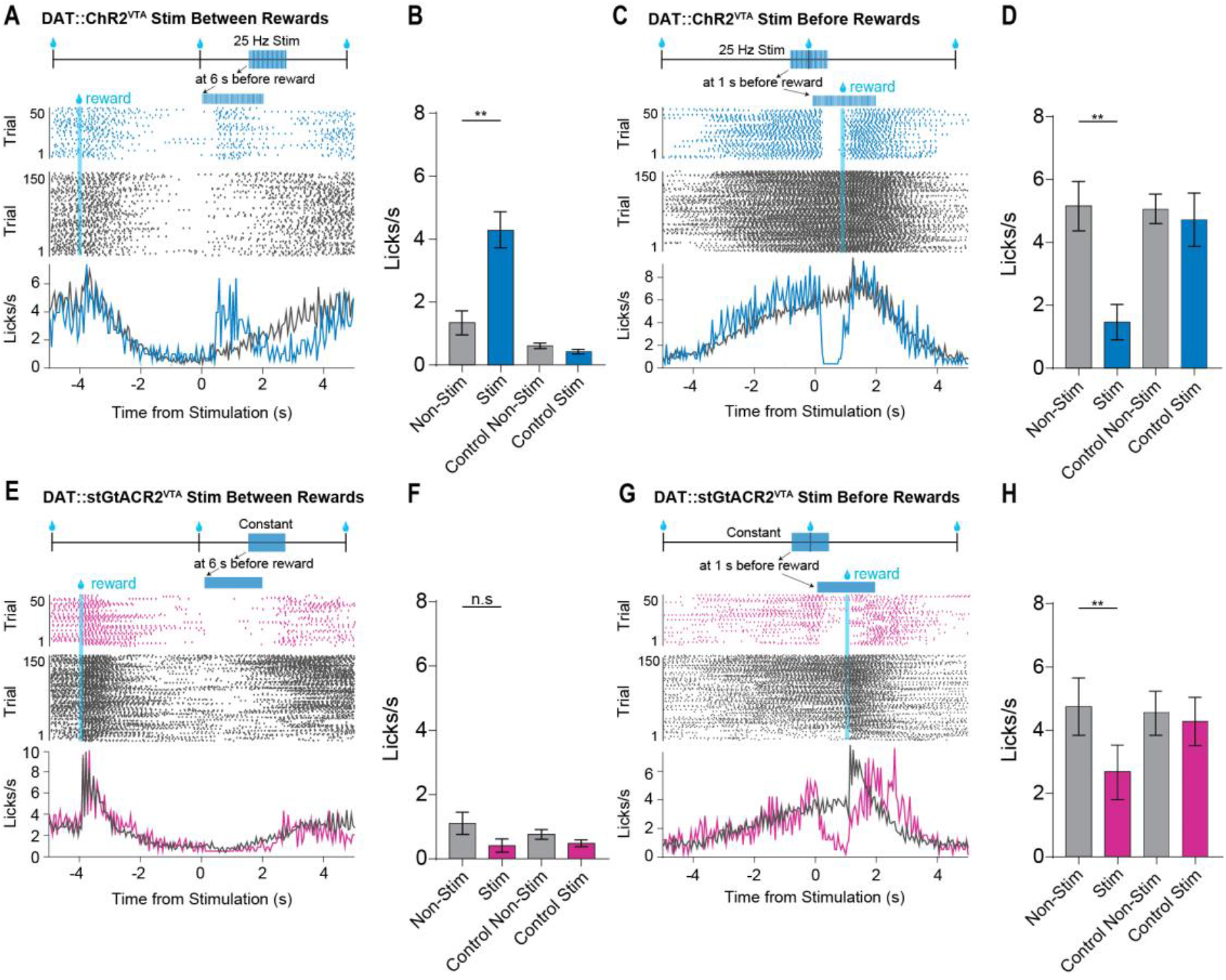
Optogenetic manipulation of VTA DA neurons initiates and disrupts anticipatory licking depending on the timing of stimulation. **A)** *Top*: Diagram of DAT::ChR2_VTA_ excitation at 6 seconds before reward delivery. *Bottom*: Optogenetic excitation produces anticipatory licking. Representative example of a licking peri-event raster plots showing licking during non-stimulation trials (grey) and stimulation trials (blue). Traces reflect the mean lick rate calculated from the raster above. **B)** The lick rate significantly increased during stimulation trials for DAT::ChR2_VTA_ (*n* = 5) animals compared to non-stimulation trials and control groups (*n* = 6; two-way mixed ANOVA, Interaction: *F*_(1,9)_ = 15.45, *p* < 0.01). **C)** *Top*: Illustration of DAT::ChR2_VTA_ excitation 1 second before reward. *Bottom*: Optogenetic excitation disrupted anticipatory licking. Representative example of licking peri-event raster plot during non-stimulation trials (black) and stimulation trials (blue) for representative animal, as well as average traces of lick rate. **D)** The rate of licking prior to reward delivery was significantly decreased during stimulation trials for DAT::ChR2_VTA_ (*n* = 5) animals compared to non-stimulation trials and control groups (*n* = 6; two-way mixed ANOVA, Interaction: *F*_(1,9)_ = 15.14*,p* < 0.01). **E)** *Top*: Illustration of DAT::stGtACR2_VTA_ inhibition 6 seconds before reward. *Bottom*: Optogenetic inhibition slightly disrupted anticipatory licking. Representative peri-event raster plots of licking during non-stimulation trials (black) and stimulation trials (maroon), with mean lick rate plotted below. F) The rate of licking did not significantly decrease during stimulation trials compared to non-stimulation trials for DAT::stGtACR2_VTA_ (*n* = 4) animals and control groups *(*n* =* 6; two-way mixed ANOVA, Interaction:, *F*_(1,8)_ = 2.36, *p* > 0.05), though this was due to a floor effect. G) *Top*: Illustration of DAT::stGtACR2_VTA_ inhibition 1 second before reward. *Bottom*: Optogenetic inhibition 1 second prior to reward delivery significantly disrupted anticipatory licking. Peri-event raster plots of licking and average traces of lick rate during non-stimulation trials (black) and stimulation trials (maroon) for representative animal. H) Lick rate significantly decreased during stimulation trials for DAT::stGtACR2_VTA_ (*n* = 4) animals compared to non-stimulation trials and control groups (*n* = 6; two-way mixed ANOVA, Interaction:, *F*_(1,8)_ = 20.13, *p* < 0.01). *p* values are corrected for multiple comparisons using Dunnett’s multiple comparison test. * *p* < 0.05; *\*\*p* < 0.01. Error bar indicates SEM.

## Discussion

Collectively, our results demonstrate that VTA DA controls the exertion of force in motivated behavior. Strikingly, we discovered a precise quantitative relationship between the activity of VTA DA neurons and the impulse vector. Distinct populations of DA neurons are associated with different components of the impulse vector. The FF and FB populations are responsible for sudden force changes needed when salient stimuli are encountered. In contrast, the SF population is responsible for slower and more gradual force generation during anticipatory approach behavior. In normal behavior, all three populations are involved, but their relative contributions depend on the force requirements of the movements needed to move towards or away from salient stimuli. Thus the magnitude and direction of the movement depends on the intensity and motivational relevance of the stimuli encountered.

In freely moving animals, it has been previously shown that DA neurons in the nearby substantia nigra pars compacta (SNc) are highly correlated with the vector components of velocity and/or acceleration [33], just like their target neurons in the striatum [34]. Optogenetic experiments also demonstrated that selective stimulation of SNc DA neurons can directly initiate movements [33], and optogenetic inhibition of SNc DA neurons can retard the initiation of movements [16]. In agreement with these observations, we showed that VTA DA neurons are also critical for the initiation of movements.

Our optogenetic experiments showed that every pulse of light exciting VTA DA neurons produced a proportional increase in impulse, demonstrating a causal role of these neurons in force generation and a linear relationship between DA activity and impulse (Figure 4). Impulse, or change in momentum, is necessary for the initiation of any movement.

According to the impulse-momentum theorem:

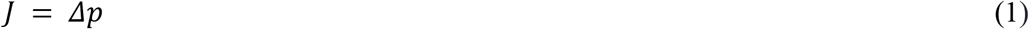

where *J* is impulse and *p* is momentum. Since momentum is defined as:

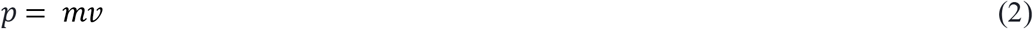

where *m* is mass and *v* is velocity, the relationship between impulse and velocity is:

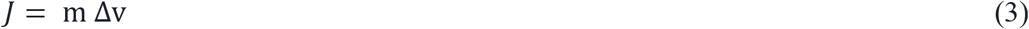

From equation (3), and assuming a constant mass, impulse simply changes velocity. Thus, it is easy to see how DA activity could be correlated with velocity and acceleration in freely moving animals [33, 35, 36], while representing impulse or change in momentum in our head-fixed preparation.

Furthermore, in naïve mice stimulation of VTA DA neurons did not generate movements, an observation suggesting that the role of these neurons in movement initiation is not unconditional, as would be the case for motor neurons. Instead, the effect of VTA DA neurons depends on the motivational context. This could explain why others have not been able to elicit movement by mimicking phasic DA activity [28]. This is in accord with the idea that DA, as a neuromodulator, primarily adjusts the gain striatal medium spiny projection neurons[37]. Striatal neurons receiving DA projections require glutamatergic drive as well as DA to be activated. The VTA DA signal alone may not be sufficient to generate movements, but instead regulates the gain of context-specific commands for motivated behaviors [38, 39].

### VTA DA regulates anticipatory licking

Another novel finding from our study is that stimulation of VTA DA neurons not only generated force but also licking behavior, even though their firing rates represented impulse rather than licking (Figure 5). When stimulated in the absence of any anticipatory licking, stimulation simply induces licking. This finding suggests that the underlying mechanisms for force generation are upstream of the licking initiation and play a key role in generating licking behavior. When stimulated during anticipatory licking, however, stimulation can actually have a disruptive effect on performance. Thus, VTA DA may induce anticipatory lick generation in the appropriate motivational context, in accord with the gain control mechanism. In contrast, inhibition of DA neurons generally suppressed anticipatory licking. Surprisingly, neither excitation nor inhibition significantly affected consummatory licking, so that the effect of DA manipulation appeared to be limited to anticipatory reward seeking behavior only.

Because we also showed that DA activity is not correlated with licking (Figure 5), these results also support the hypothesis that VTA DA may regulate the gain of context-dependent behavioral sequences. This type of regulation is manifested in the force generated. This is consistent with extensive past research implicating VTA dopamine in effort, wanting or vigor related to motivationally relevant stimuli[7, 40].

It has recently been reported that VTA GABA neurons, which are directly connected with the DA neurons, can represent and command head angles about orthogonal axes of rotation[29]. Given the interaction between VTA GABA and DA, DA could contribute to head angle commands by providing the requisite amount of torque necessary to achieve a change in head position (i.e. angular velocity) in order to move toward or away from motivationally relevant stimuli. It should be noted, however, the force measured in our study cannot be used to predict actual kinematics in freely moving animals, because the force sensors do not accurately measure torque due to the geometry of the body in the head-fixed animal and the positioning of the limbs and associated mechanical advantage. In addition, depending on the mass of the body, some minimum amount of force must be generated for overt movement.

### Reward Prediction Error

Although our findings do not rule out a role of DA in learning, they are inconsistent with the popular RPE hypothesis of DA function[4] for the following reasons: 1) All three populations of VTA DA neurons increased their firing rate regardless of the motivational valence of the stimuli involved (reward or air puff, Figure 3). 2) In our paradigm, rewards are fully predicted, yet post-reward DA bursts are always observed. This could be due to uncertainty as reflected in timing precision, but we found that variability in the timing of anticipatory licking, a measure of how well the reward is predicted, is unrelated to the phasic burst response of DA neurons after reward delivery (Figure 5). 3) Each population of DA neurons shows a unique pattern after reward delivery, corresponding to distinct components of the impulse vector and exhibited clear and opposite changes in their firing rates depending on the direction of force (Figures 3–4). 4) Optogenetically mimicking phasic RPE bursts generated net forward force, and optogenetic inhibition could result in backward force generation (Figure 6).

More generally, the RPE hypothesis only predicts increases or decreases of DA activity based on the magnitude of the prediction error. It is not a vector quantity and contains no spatial or directional component, despite previous attempts to reconcile learning and performance in such models [6, 7, 41]. Previous experiments in support of the RPE hypothesis neglected confounding variables associated with movements. This is especially true of head-fixed experiments, where overt movements are subtle and difficult to detect. Nevertheless, just because animals are prevented from expressing clear movements does not mean that they do not attempt to move. Only with careful behavioral measures, such as the sensitive force sensors used here, can such subtle behaviors be quantified.

Recent work by Coddington and Dudman also found that VTA DA neurons can both be inhibited or excited during the initiation of movement [28]. Although they attempted to quantify movement in a head-fixed preparation, their behavioral measures lacked spatial resolution. For example, the accelerometer baskets they used could not distinguish between forward and backward movements. Our more sensitive behavioral measures revealed that there are discrete VTA DA populations based on the direction and amplitude of head movement. Furthermore, there is often a pronounced, but very brief, backward movement during reward delivery which coincides with the phasic burst. While their proposed model also attempts to reconcile movement initiation effects and RPE, it lacks vector quantities, and leaves action vaguely defined, a major limitation in all reinforcement learning models. Consequently, it cannot explain the present results.

Our findings indicate that phasic dopamine bursts signal a sudden change in force associated with rapid but subtle movement in response to salient stimuli. The direction of the movement is determined by the force vector. Variable patterns and directions of movement produced in different experimental set ups could explain why conflicting results have been found for the ‘encoding’ of aversive and rewarding events in projection-specific populations of VTA DA neurons [24, 42]. It would be critical for future studies examining the functional heterogeneity in DA neurons to carefully quantify movements in different directions, which can vary with standard manipulations intended to vary reward magnitude, probability, and valence[15, 43].

More recent work has claimed that VTA DA neurons can multiplex many different variables, including reward and movement [11, 12]. While this interpretation may appear on the surface to explain the greatest number of previous findings, it is also challenged by our results. First, the statistical methods used in such studies allow them to describe nearly any type of neural activity as multiplexing any number of behavioral variables. In reality, there is no clear relationship between the neural activity and any of the variables measured, as indicated by very low correlations between behavioral variables and DA activity, probably due to the technical limitations of conventional measures as explained above. Without an explicit model of how the signals can be de-multiplexed, this interpretation is not even falsifiable. More importantly, these studies did not measure the key variable, namely the impulse vector, that actually accounts for most of the variance in the activity of DA neurons (Figure 4). In our study, we were also able to establish a direct causal connection with our optogenetic experiments. Thus, task-relevant variables used in models that purport to predict DA activity may in fact be correlated with forces exerted by the animal or vector components of kinematic variables [33, 34]. Previous studies may have omitted a universal property of the animal’s behavior that co-vary with all the variables selected by the experimenter for analysis.

### Impulse and Motivation

Our results are more consistent with the idea that VTA DA plays a key role in motivation by energizing behavior. It has been proposed that VTA DA represents incentive salience of motivationally relevant stimuli, and how much vigor or effort is being expended towards these stimuli [6, 7, 44, 45]. In addition, there is an extensive literature linking mesolimbic activity and locomotion[5, 38, 39, 46]. One limitation of these proposals is that, like the RPE hypothesis, they also lack vector quantities. Vigor, for example, is not a vector quantity with information about direction, and cannot generate specific behavioral trajectories. Our findings confirm and significantly extend such work. We show that the impulse vector explains the activity of most VTA DA neurons, and suggests a simple, quantitative, and easily falsifiable hypothesis of VTA DA function.

In summary, our results indicate that VTA DA neurons regulate anticipatory behavior by controlling the force required to move in specific directions and initiate the appropriate behavioral sequence. We show that the signals carried by the VTA DA neurons represent vector quantities that can be used to predict both the direction and amplitude of forces exerted over time, which is a significant advance towards predicting continuously generated behavior.

## Acknowledgments

We would like to thank Dr. Fengxia Allen, Dr. Guozhong Yu, and Murray Wickwire for their technical assistance. We would like to thank Dr. Joseph Barter for assistance with the head-fixation apparatus, as well as helpful comments and discussions.

## Funding

This work was supported by NIH grants NS094754, DA040701, and MH112883 to HHY.

## Author contributions

R.N.H., K.I.B., and H.H.Y. conceptualized and designed studies. R.N.H and K.I.B., performed surgeries and optogenetic experiments. R.N.H. performed *in vivo* electrophysiological experiments. A.D.F. performed behavioral experiments. N.M.K performed *in vitro* electrophysiological recordings. G.D.R.W. and R.N.H performed immunohistochemistry and confocal imaging. R.N.H., E.A.P, and K.I.B wrote data collection and analysis code. E.A.P performed data analysis. R.N.H., K.I.B and H.H.Y. drafted and edited the manuscript. All authors have read and approved the manuscript.

## Competing interests

The authors report no financial interests or potential conflicts of interest.

## Supplemental Materials

STAR Methods

Figures S1-S7

Movies S1-S2

