## Supplemental information for "Ventral tegmental dopamine neurons control the impulse vector during motivated behavior"

**This PDF file includes:**

STAR Methods
Figs. S1 to S7
Captions for Movies S1 & S2

**STAR Methods**

**Key Resources Table**

| ***REAGENT or RESOURCE*** | ***SOURCE*** | ***IDENTIFIER*** |
|---|---|---|
| ***Antibodies*** | | |
| Goat polyclonal anti-Rabbit Alexa Fluor 488 | Abcam | Cat# ab150077; RRID: AB_2630356 |
| Goat polyclonal anti-Rabbit Alexa Fluor 594 | Abcam | Cat# ab150080; RRID: AB_2650602 |
| Rabbit polyclonal anti-Dopamine Transporter | Abcam | Cat# ab111468; RRID: AB_11155293 |
| Rabbit polyclonal anti-Tyrosine Hydroxylase | Millipore | Cat# 657012; RRID: AB_566341 |
| ***Bacterial and Virus Strains*** | | |
| rAAV5-EF1α-DIO-eYFP | Duke Vector Core | N/A |
| rAAV5-EF1α-DIO-hChR2(H134R)-eYFP | Duke Vector Core | N/A |
| AAV1-hSyn-SIO-stGtACR2-FusionRed | Addgene | Cat# 105677-AAV1 |
| ***Experimental Models: Organisms/Strains*** | | |
| Mouse: DAT-ires-Cre: 1$^{tm2(cre)Lowl}$/J | Jackson Laboratory | Mouse Strain: 006660 |
| Mouse: C57BL/6J | Jackson Laboratory | Mouse Strain: 000664 |
| Mouse: Ai32: B6;129S-*Gt(ROSA)26Sor*$^{tm32(CAG-COP4*H134R/EYFP)Hze}$/J | Jackson Laboratory | Mouse Strain: 012569 |

| | | |
|---|---|---|
| ***Software and Algorithms*** | | |
| MATLAB 2016b | MathWorks | https://www.mathworks.com/products/new_products/release2015b.html |
| Python 2.7 | Anaconda | https://www.anaconda.com/download/?lang=en-us |
| Offline Sorter 3.0 | Plexon | https://plexon.com/products/offline-sorter/ |
| NeuroExplorer 4.0 | Nex Technologies | http://www.neuroexplorer.com/downloadspage/ |
| Cortex 5.0 | MotionAnalysis | http://ftp.motionanalysis.com/html/industrial/cortex.html |
| GraphPad Prism 8 | GraphPad | https://www.graphpad.com/scientific-software/prism/ |
| ***Other*** | | |
| LED Driver | Thorlabs | LEDD1B |
| DAPI Fluoromount-G | Southern Biotech | Cat# 0100-20 |

**Experimental Model and Subject Details**

All experimental procedures were approved by the Animal Care and Use Committee at Duke University. 13 *DAT-ires-cre* mice, 4 *DAT-Cre + Ai32* mice and 6 *wild-type* male and female mice were used for experiments with virally delivered opsins (Jackson Labs, Bar Harbor, ME). For experiments involving endogenously expressed opsins, Ai32 mice were crossed with DAT-Cre animals (both heterozygous and homozygous *DAT-ires-Cre* mice). All mice were aged between 2-8 months old. Mice were housed on a 12:12 light cycle, with tests occurring in the light phase. All mice were housed in groups of 3-4 animals per cage. For experiments, mice were put on water restriction and maintained at 85-90% of their initial body weights. Animals received free access to water for approximately 30 minutes following daily experimental sessions.

### Method Details

***Viral Constructs***. rAAV5.EF1α.DIO.hChR2(H134R) was obtained from the Duke University Vector Core. pAAV-hSyn1-SIO-stGtACR2-FusionRed was purchased from Addgene. pAAV_hSyn1-SIO-stGtACR2-FusionRed was from Ofer Yizhar (Addgene viral prep # 105677-AAV1; http://n2t.net/addgene:105677; RRID:Addgene_105677).

***Surgery.*** Mice were initially anesthetized with 2.0 to 3.0% isoflurane before being placed into a stereotactic frame (David Kopf Instruments, Tujunga, CA) and were maintained at 1.0 to 1.5 % for surgical procedures. For experiments involving viral injections, 200-300 nL of DIO-ChR2, SIO-StGtaCR2, or DIO-eYFP were bilaterally injected at a rate of 1 nL/s into the VTA (AP: 3.2 – 3.4 mm relative to bregma, ML: 0.4 – 0.6 mm relative to bregma, DV: 4.0 – 4.4 mm relative to brain surface) using a microinjector (Nanoject 3000, Drummond Scientific). The injection pipette was left to sit for 3-5 minutes immediately after the injection to allow absorption of the virus and prevent leakage. Custom-made optic fibers (5 - 6 mm length below ferrule, >80% transmittance, 105 μm core diameter) were implanted at a fifteen-degree angle above the middle of the VTA = (AP: 3.2 - 3.4 mm, ML: 1.6 mm, DV: 3.8 mm). For electrophysiological recordings, drivable electrodes were placed just above the VTA (AP: 3.2 - 3.4, ML: 0.5 mm, DV: -3.8 mm) and 16-channel recording electrodes were lowered into the VTA (AP: 3.2 – 3.4 mm, ML: 0.5 mm, DV: -4.0 - 4.4 mm). For optotagging experiments, an optic fiber was attached to the electrodes. Fibers and electrodes were secured to the skull using screws and dental acrylic and all mice were fitted with a steel head implant for head fixation. All animals were allowed to recover for two weeks before beginning training on the fixed-time task.

***Head-fixed behavioral setup.*** We created a customized head-fixation device for measuring forces exerted by the animals during behavioral testing and stimulation. Animals' head implants were clamped to a frame that was suspended on a set of 3 load-cells (RB-Phl-203, RobotShop.com) that were arranged orthogonally to detect force changes in three dimensions (forwards-backwards, up-down, and left-right). Animals stood on an enclosed platform that incorporated two additional load-cells that detected downward forces exerted by the left and right feet. Load cells translate small mechanical distortions caused by an applied load into a voltage signal. We amplified the signal using an INA125P (Texas Instruments) in a standard circuit configuration (http://www.mechtechplace.net/mech-tech-electronics/building-a-low-cost-strain-gage-load-cell-amplifier/). A metal spout attached to a reservoir containing a 10% sucrose solution was placed close to the animals' mouth. Sucrose delivery was gravity fed and controlled by the opening of a solenoid valve (161T010, NResearch, NJ). A capacitance-touch sensor (MPR121, AdaFruit.com) clamped to the metal spout was used to detect individual licking events. All continuous load cell voltages were recorded at 1 kHz along with timestamps corresponding to reward, laser train delivery times, individual licks, and electrophysiological data (see below) using a Blackrock Cerebrus recording system (Blackrock Microsystems) for offline analysis.

***Behavioral task.*** Water-deprived animals were first habituated to the head fixation apparatus and trained to receive water rewards delivered manually by the experimenters. Once reliable licking was observed, animals were switched to the fixed-time task, in which mice receive a 5 uL drop of 10% sucrose solution every 10 seconds. Reward delivery was not contingent upon animals' behavior, so they received the reward regardless of their performance. On a subset of recording sessions (n = 6), air puffs were delivered during the interval task. The air puffs could occur randomly any time during the interval between 1 and 9 seconds with a probability of 15% for a given trial. The task was controlled using Matlab (version 2018b, Mathworks, MA) commanding digital outputs through a data acquisition system (USB6001, National Instruments, NJ).

***Wireless in Vivo Electrophysiology.*** Both fixed and drivable 16-channel electrode arrays or were used (Innovative Neurophysiology, Inc.). Drivable electrodes were single-drive movable micro-bundles of tungsten electrodes (1 x 16; 23 μm diameter) placed within a guide cannula. Fixed arrays were composed of tungsten electrodes in a 4 x 4 array (35 μm diameter, 150 μm spacing, 5 mm length). Electrophysiological data were recorded via a miniaturized wireless head stage (Triangle Biosystems) that communicated with a data acquisition system (Blackrock Microsystems). Both analog and digital bandpass filters were applied to the electrophysiological data (analog highpass 1st order butterworth filter at 0.3 Hz, analog lowpass 3rdorder butterworth filter at 7.5 kHz, digital highpass 4th order butterworth filter at 250 Hz). Filtered data were then sorted offline using OfflineSorter (Plexon). All raster plots of spiking activity and force signals were generated using NeuroExplorer (Nex Technologies) using 50 ms timebins and smoothed using a Gaussian filter with a standard deviation of 3 bins. A 3:1 signal-to-noise ratio, and an

800 μs or greater refractory period were required for the neural data to be used for analysis. For driveable electrodes, the electrodes were lowered after each session by 50 μm. In order to be considered a different unit within the same channel, the waveform had to be significantly different from the previously recorded unit as determined by OfflineSorter. For optotagging experiments, optic fibers were epoxied to driveable electrodes.

**Histology**: Mice were transcardially perfused with 0.1M phosphate buffered saline (PBS) followed by 4% paraformaldehyde (PFA) in order to confirm viral expression as well as optic fiber and electrode placement. To confirm placement, brains were stored in 4% PFA with 30% sucrose for 72 hrs. Tissue was then post-fixed for 24 hours in 30% sucrose before cryostat sectioning (Leica CM1850) at 60 μm. Fiber and electrode implantation sites were then verified. To confirm eYFP and FusionRed expression in *DAT+* cells in the VTA of *DAT-ires-Cre* and *DAT + Ai32* transgenic mice, sections were rinsed in 0.1M PBS for 20 min before being placed in a PBS-based blocking solution. The solution contained 5% goat serum and 0.1% Triton X-100 and was allowed to sit at room temperature for 1 hr. Sections were then incubated with a primary antibody (polyclonal rabbit anti-TH 1:500 dilution, ThermoFisher, catalog no. P21962; polyclonal chicken anti-EGFP, 1:500 dilution, Abcam, catalog no. ab13970) in blocking solution overnight at 4 °C. Sections were then rinsed in PBS for 20 min before being placed in a blocking solution with secondary antibody used to visualize *DAT* neurons in the VTA (goat anti-rabbit Alexa Fluor 594, 1:1000 dilution, Abcam, catalog no. ab150080; goat anti-chicken Alexa Fluor 488, 1:1000 dilution, Life Technologies, catalog no. A11039) for 1 hr at room temperature. Sections were mounted and immediately coverslipped with Fluoromount G with DAPI medium (Electron Microscopy Sciences; catalog no. 17984-24). Placement was validated using an Axio

Imager.V16 upright microscope (Zeiss) and fluorescent images were acquired and stitched using a Z780 inverted microscope (Zeiss).

**Whole-cell patch clamp recording:** For whole-cell patch-clamp recordings, 3 *DAT-Cre* animals were used. StGtACR2 was injected into the VTA and the mice were sacrificed 8-12 weeks after the injection. The brain was removed quickly and left in ice-cold solution bubbled with 95% O2-5% CO2 containing the following (in mM): 194 sucrose, 30 NaCl, 2.5 KCl, 1 MgCl2, 26 NaHCO3, 1.2 NaH2PO4, and 10 D-glucose. After 5 minutes, 250 µm coronal slices were cut and then placed in 35.5°C oxygenated artificial cerebrospinal fluid (aCSF) solution containing the following (in mM): 124 NaCl, 2.5 KCl, 2 CaCl2, 1 MgCl2, 26 NaHCO3, 1.2 NaH2PO4, and 10 D-glucose. After 30 minutes, the slices were left in aCSF at ~22 -23°C for at least 30 min before recording. Following recovery, whole-cell patch clamp recordings were performed in current clamp mode with continuous perfusion of aCSF at 29-30°C. The internal solution contained (in mM) 150 potassium gluconate, 2 MgCl2, 1.1 EGTA, 10 HEPES, 3 sodium ATP, and 0.2 sodium GTP.

To measure the inhibition by light stimulation, slices were injected with current that was adjusted to evoke action potentials (30 – 150 pA). Current was delivered for 2 seconds; and 500 ms after the start of current injection slices were stimulated with 470-nm light from an LED (Thor Labs). Both constant (1 s) and pulsed stimulation (5 ms pulses at 10, 25 and 50 Hz; MASTER-8) was delivered to the entire ×40 field with an LED current driver (Thor Labs). Power density was ~2 $mW/mm^2$. Action potentials were recorded for 500 ms after light stimulation. All recordings were performed with a MultiClamp 700B amplifier (Molecular Device). Signals were filtered at 10 kHz and digitized at 20 kHz with a Digidata 1440A digitizer

(Molecular Devices). The inhibition ratio was measured by comparing firing rates from before and after light stimulation.

**Optogenetic experiments**:

*Stimulation schedule*: Animals were well trained before beginning optogenetic manipulations. Optogenetic stimulation sessions were identical to electrophysiological sessions described above, except for the delivery of light on pseudo-randomly determined trials (see diagram, Figure S6A). After 2 consecutive reward deliveries, the program would enter into a choice point. The probability of the interval following the 3rd reward containing a laser presentation was 50%. Whether laser was presented or not, the consecutive reward counter was reset to 0 after the delivery of the 3rd reward. The overall probability of receiving laser stimulation for a given training session was approximately 15%. For experiments with ChR2, optical power was measured at 8 mW at the end of the fiber that connected to the optical implants. For experiments with stGtaChR2, power was capped at 5 mW.

**Quantification and Statistical Analyses**

All analyses were performed using Matlab, Python, NeuroExplorer, and Graphpad Prism. All statistical analyses were performed in MATLAB and GraphPad Prism. A power analysis was not conducted to determine sample size *a priori.*

***Force conversions***: To calculate the forces exerted by animals on the load-cells, we calibrated each load-cell circuit by determining a linear conversion factor (expressed in Newtons per Volt) between the load cell circuit output and known masses placed on the sensor. Force was determined by multiplying the recorded voltage signal by the conversion factor to obtain the force in Newtons exerted by the animals on each load cell.

***Detection of movement initiation*:** We applied thresholds to the forward/backward force signal to detect forward and backward movement events. Forward movements were defined as continuous events exceeding 150 mN lasting longer than 250 ms and separated by at least 100ms. Backward movements were defined as events lower than –300 mN lasting longer than 100 ms and separated by at least 100 ms.

***Optical tagging and unit classification*:** Tagging of dopamine neurons was performed during the performance of the task with laser stimulation occurring at –6 seconds prior to reward. Neural spiking data was recorded simultaneously with timestamps for optical stimulation pulses. Neural data and optic stimulation timestamps were imported into NeuroExplorer and aligned to the start of stimulation. Neurons were classified as tagged DA neurons if the first evoked action potential had a latency of less than 6 ms and resulted in a waveform identical to the spontaneously-occurring waveform for that unit. In order to classify non-tagged neurons as DA, optically tagged and non-tagged waveforms were exported from Plexon into Matlab, where the valley full-width half max (FWHM) was calculated using a custom Matlab script. The average FWHM and firing rate were then computed. The majority of the optically tagged neurons had a valley FWHM of at least 500 μs and a firing rate below 10 Hz. The remaining non-tagged DA neurons were classified as VTA DA neurons if their firing rate was below 10 Hz and their valley FWHM was over 500 μs.

***Functional classification of DA neurons*:** Neurons were initially manually observed as either a fast-forward (FF), fast-backward (FB), or slow-forward (SF) DA neurons based on their activity during the time of reward, forward and backward movement. To further confirm and validate distinct functional VTA DA populations, an unbiased clustering algorithm (Gaussian mixture model [GMM]) was used. Neural data from units classified as DA were aligned to reward, forward movement and backward movement in Neuroexplorer, and then exported to Microsoft Excel. Data were then concatenated into one response profile and analyzed using Python 3 and the sci-kit learn package. Data was smoothed with a Guassian filter (sigma=2), and then scaled (MinMaxScaler). The scaled response profiles were whitened, and a PCA decomposition (3 components) was performed on the data. The resulting principal components were clustered using a GMM (3 clusters). Over 95% of neurons initially classified as an FF, FB, or SF neuron fell within the same cluster using the GMM.

***Analysis of behavioral and neural data*****:** To examine the relationship between neural activity and force generated over time (impulse), we first identified all forward and backward movement events occurring in a recording session (see threshold criteria above). We compared spiking activity of FF and SF neurons to impulse of forward movements, and spiking activity of FB neurons to impulse of backward movements. Impulse was estimated using the *trapz* function in Matlab as the area under the curve of F/B force over the duration of each movement. We also recorded the peak force for each movement and counted the number of spikes recorded for each DA neuron in the same time interval. Thus, for each neuron and each movement event, we obtained a single number for impulse, a single number for peak of the force signal, and a single number of spikes. For each neuron, we correlated all spike count values with the corresponding impulse values, and obtained an average impulse per spike count for each cell. These values were averaged across neurons from each functional class. In order to compare across different neurons, we plotted impulse as a function of spike number up to 7 spikes (7 was the maximum spike number that all neurons have in common during a movement). To account for the fact that neural activity often preceded force generation, for each neuron we adjusted the time window used to count spikes associated with each movement by the lag estimated from the cross-correlation with force over the entire session (see method for cross-correlation below).

***Analysis of force in optogenetic experiments*:** Force signals were aligned to the onset of laser delivery in NeuroExplorer and binned at 50 ms. The mean peri-event force signal across all laser presentation trials in the forward/backward direction was exported to Matlab where we estimated the maximum value and also found the area under the curve using the *trapz* function. This resulted in one value for each animal for each laser stimulation condition. Trials were excluded from this average if laser delivery did not evoke any movement. These values were averaged together before regression analysis with pulse number. For estimation of force generated in control animals, we obtained the average duration of the movements evoked by a given stimulation parameter. We then determined the maximum force and measured impulse of the mean force signals that were recorded over the same durations following laser activation in control experiments. To quantify negative impulses that resulted from inhibition of VTA DA neurons, we superimposed the mean force signals generated during trials with and without laser, and measured the area between the two curves. This impulse measure was determined for the first second of laser presentation which coincided with the anticipatory phase of the interval. Impulse resulting from inhibition rebound was determined in the same manner as with ChR2 experiments described above.

***Analysis of Forward Force Movements with No Lick Bouts:*** In order to determine if VTA DA neural activity is still correlated with force in the absence of lick bouts, identified non-rewarded forward and backward movement events that occurred outside of periods of licking. Lick bouts consisted of licking events with a maximum inter-lick-interval of 0.11 s and had to contain more than one lick. Neural data was then aligned to these filtered forward movements. All three populations showed similar relationships with F/B force during periods without any reward delivery or lick bouts being initiated (Figure S6).

***Support Vector Regression Decoder:*** Support vector regression was implemented via the *scikit-learn* Python package to fit the continuous forces recorded from each load cell (forward/back, up/down, side/side) using the firing rate (bin size: 50 ms) of DA neurons. As the number of neurons and the size of the training data affects decoding performance, we limited our analysis to sessions that contained at least 6 DA neurons (63% of datasets; maximum number of DA neurons was 20). Prior to fitting the model, the neural data was z-scored, and the force was zero-centered. We then convolved the force data with a Gaussian filter (width: 5 bins). For each dataset, the model was trained on the first 60% of the data, and performance was evaluated on a continuous set of held-out data (15%).

***Correlation Analyses:***

*Correlation Analyses Across Entire Behavioral Session*: For each animal, neural data and continuously monitored F/B force or lick rate for the entire recording session were constructed using 10 ms time bins in NeuroExplorer and exported to MATLAB. A custom matlab script was then used to sort neural firing rate variables and the corresponding force signal or licking rate according to firing rate magnitude. Neural activity was binned into twenty bins, and the mean force signal or licking rate corresponding to the firing rate in each bin was calculated. The

Pearson correlation was calculated between binned neural activity and mean force signal or licking rate.

*Cross-Correlation Analyses:* A custom MATLAB script was written in order to perform cross-correlation analyses between the force measurements and the corresponding VTA DA neuronal subtype. Cross-correlations were performed in MATLAB by utilizing the in-built function *xcorr*. The behavioral variable was used as the reference time-series and the neural data as the shifted time series. Latencies were obtained by determining the lag of the maximum value of the cross-correlation for positively correlated neurons and the lag to the minimum value for anti-correlated neurons. Cross-correlations were then Z-score normalized and all data sets from all animals were averaged together to create population average plots.

**Data and Code Availability**

Raw data from the current study were not deposited into a public repository due to the large size of the data sets, but are available from the corresponding author upon request.

**Supplemental Figures**

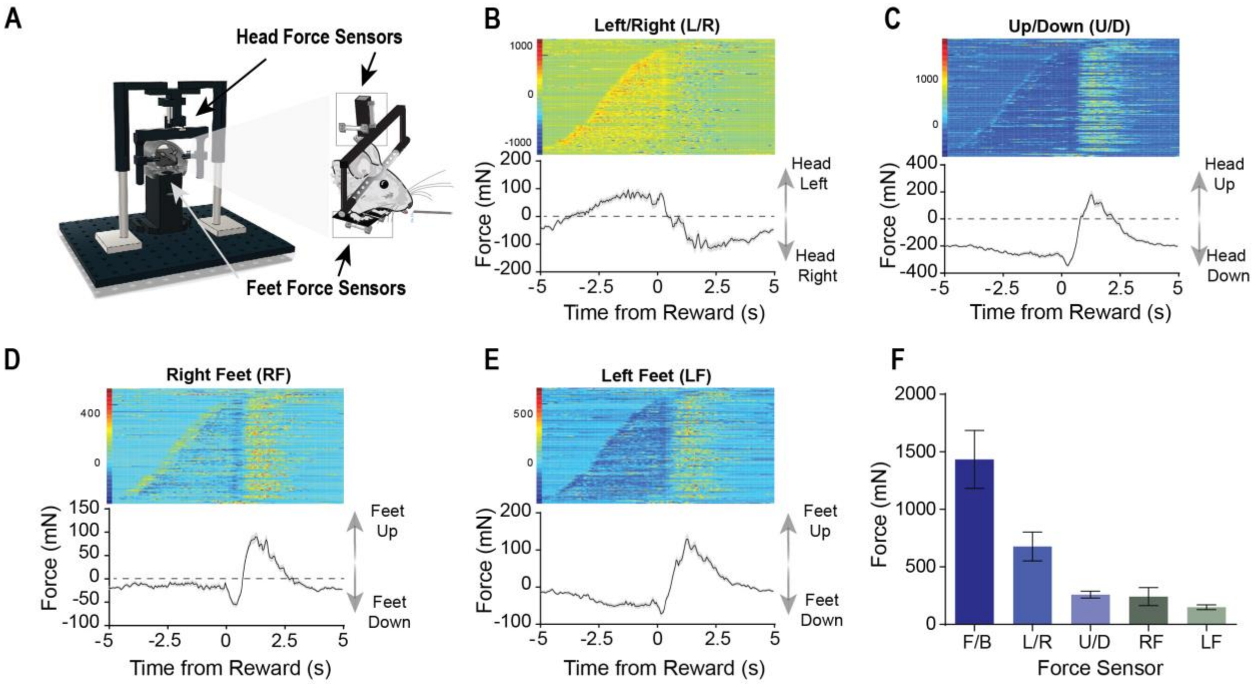

**Fig. S1. Schematic of head-fixation apparatus for studying force and examples of force sensor data**

**A**) *Left*: Schematic of novel head-fixation apparatus. *Right*: Zoomed in view of the mouse in the head-fixation apparatus with three orthogonal force sensors for the head and two for the feet. **B-E**) Representative examples of force plots for Left/Right (LR) (**B**), Up/Down (UD) (**C**), Right Feet (RF) (**D**) and Left Feet (LF) (**E**) within a single session for a single mouse (same session as shown in Figure 1). The trials were sorted based on the duration of the lick bout. **F**) The peak force exerted in the forward/backward direction was the largest and most apparent. The mice would also move left and right, but not as much. Error bars indicate SEM.

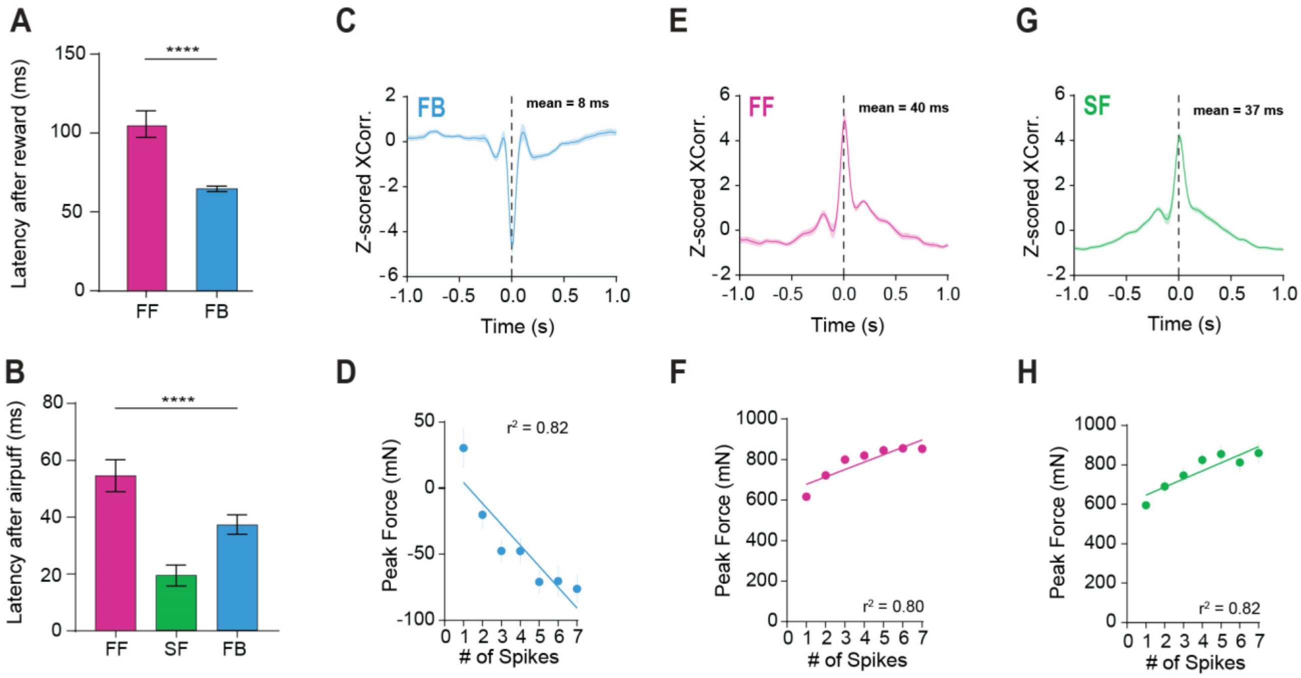

**Fig. S2. Latency analysis and correlation with peak force in each population of VTA DA neurons**

**A)** There is a significant difference between FF and FB neurons with respect to their latency of phasic bursting after reward (unpaired t-test, $t(61) = 5.46$, $p < 0.0001$). **B**) The SF population fired first after air puff, followed by the FB population, then the FF population. They were significantly different (One-way ANOVA, $F_{(2,49)} = 13.25$, $p < 0.0001$. Tukey's *post-hoc* test revealed significant differences between all three groups $p < 0.01$). **C**) Cross-correlogram for FB DA activity and F/B force (reference: F/B force). FB DA activity generally preceded F/B force. A positive value indicated the neural activity is leading the behavioral change (8 ± 14 ms). **D**) Correlation between the number of spikes of FB neuron population and the peak backward force. **E)** Cross-correlogram for FF DA activity and F/B force. FF DA activity generally preceded F/B force. (40 ± 17 ms). **F**) Correlation between the number of spikes of FF neuron population and the peak forward force. **G**) Cross-correlogram for SF DA activity and F/B force. SF DA activity generally preceded F/B force. (37 ± 14 ms). **H**) Correlation between the number of spikes of SF neuron population and the peak forward force. Error bars indicate SEM.

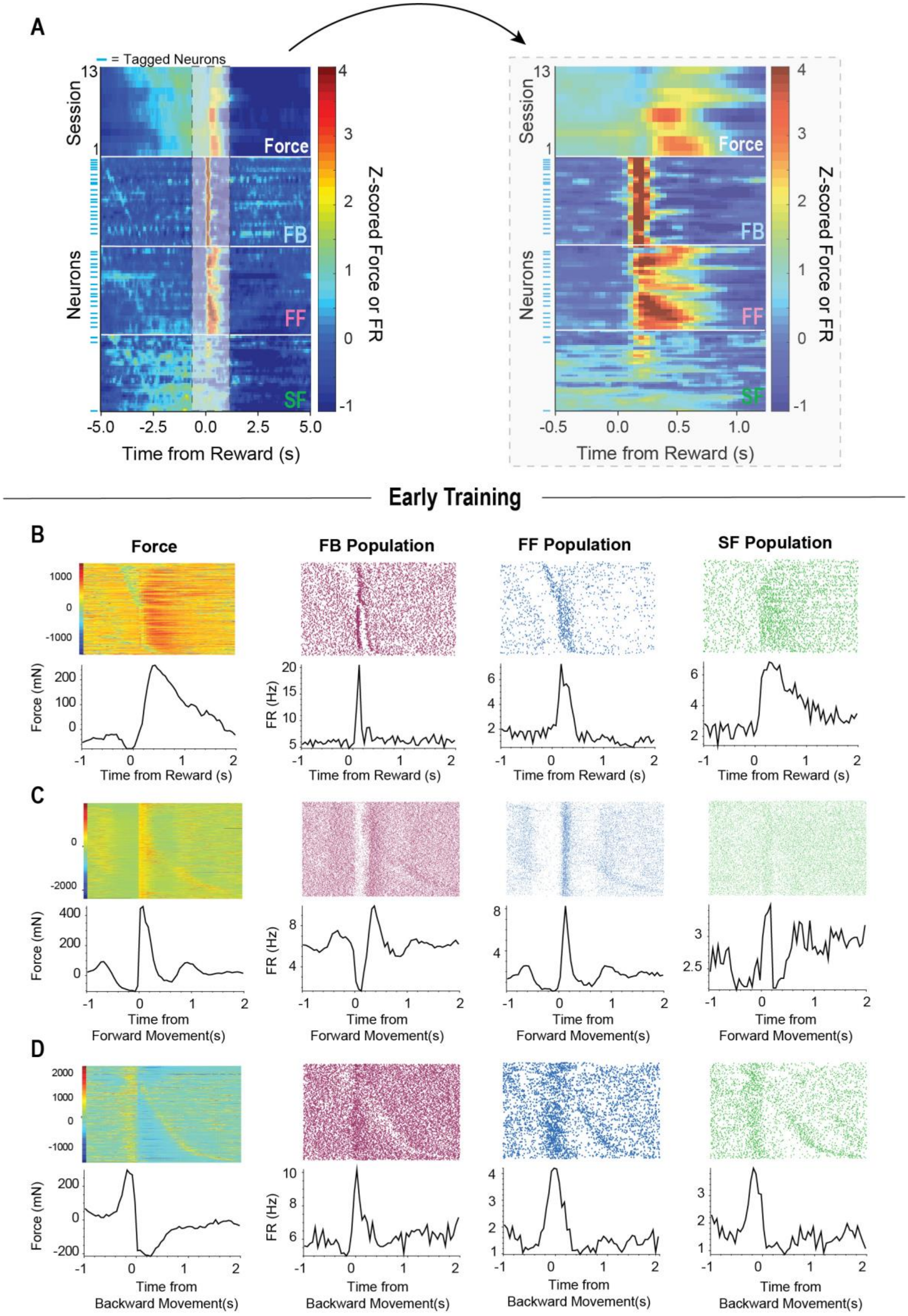
A
= Tagged Neurons
Session
Neurons
Force
FB
FF
SF
Z-scored Force or FR
Time from Reward (s)
Early Training
B
Force
FB Population
FF Population
SF Population
Force (mN)
FR (Hz)
Time from Reward (s)
C
Time from Forward Movement(s)
D
Time from Backward Movement(s)

**Fig. S3. SF population shows tonic activity while FB and FF population show more phasic activity in response to reward. The same responses are seen during early training.**

**A)** *Left:* Peri-event heat map of FB force and all neuronal populations aligned to reward. FB neurons show short phasic bursts that are concurrent with the pause in force or backward movement (burst duration < 200 ms). Positive values indicate force in the forward direction. FF neurons show long-phasic bursts (burst duration > 200 ms) that scale with the large and fast force changes after reward. SF neurons do not show phasic bursts, but scale with slow forward force changes. *Right:* Same data as *left* but zoomed in around the time of reward. Note that FB population shows a phasic burst during a brief backward movement (negative values in force). Blue ticks indicate optically tagged DA neurons. **B**) F/B force and single-unit VTA DA neurons from each population during early training aligned to reward. At this stage in training, mice do not anticipate the reward, and only move after the reward. **C**) F/B force and single-unit VTA DA neurons from each population aligned to forward movement. **D**) F/B force and single-unit VTA DA neurons from each population aligned to backward movement. Neural activity in early training sessions closely follows force changes observed during late stages of training.

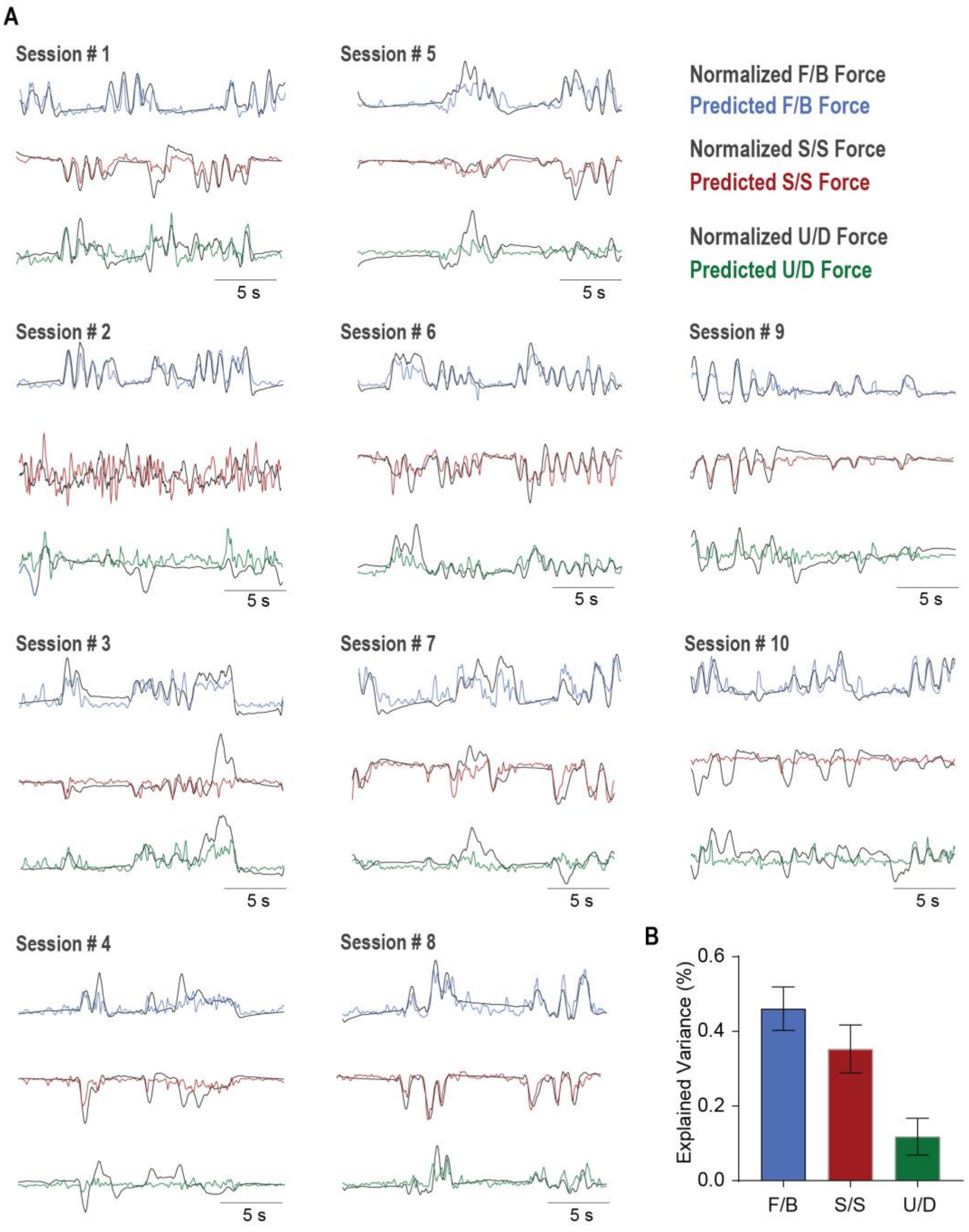

**Fig. S4. Support Vector Regression (SVR) decoder data**
**A)** 10 sessions from mice with at least 6 VTA DA neurons were decoded using each of three head force sensors (F/B, S/S, & U/D). 20 second excerpts from predicted force data are shown. **B**) Decoding performance was significantly different between F/B force, S/S and U/D force (RM one-way ANOVA, $F_{(9,18)} = 15.91$, $p < 0.0004$). Error bars indicate SEM.

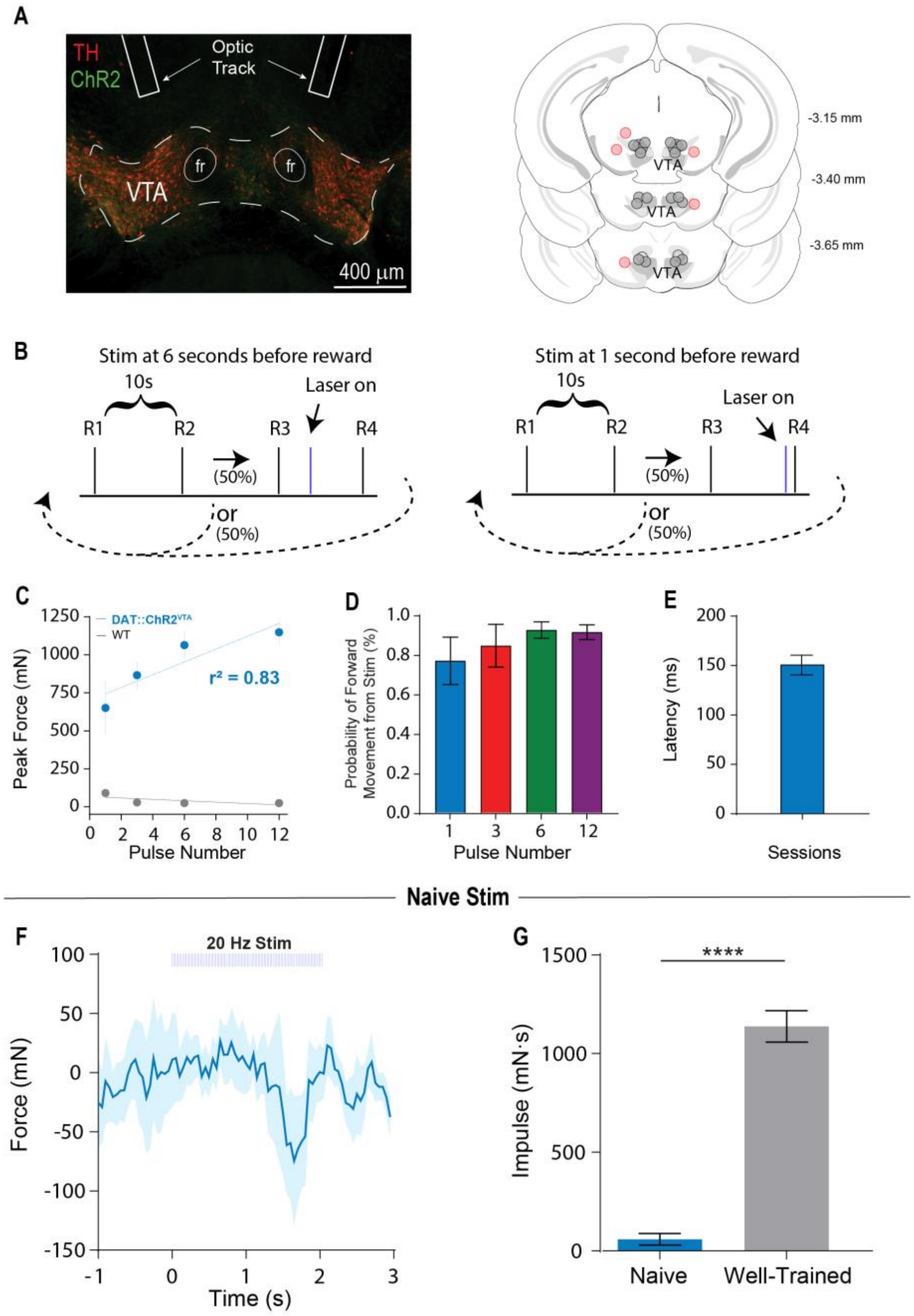

**Fig S5. Schematic of Optogenetic behavioral task design and supplemental graphs**
**A)** Histological verification of fiber placement above the VTA. Red circles denote animals that were excluded from data set due to inaccurate placement. **B**) Schematic of optogenetic stimulation schedule. **C**) A strong linear relationship (though not as strong as Impulse, $r^2 = 0.83$, $p > 0.05$) between mean peak forward force signals as a function of number of light pulses in DAT::ChR2$_{VTA}$ (n = 4) and WT animals (n = 5). **D**) Probability of forward movement was high for all number of pulses, even 1 pulse (1 pulse: 0.77 ± 0.24; 3 pulses: 0.85 ± 0.22; 6 pulses: 0.92 ± 0.08; 12 pulses: 0.91 ± 0.07). **E**) Average latency (151.50. ± 9.36) to movement onset from single pulse of light. Error bars indicate SEM. **F**) Optogenetic stimulation in naïve mice (20 Hz

for 2 seconds) did not reliably produce forward force. **G**) The impulse produced by stimulation in mice within the task was significantly greater than naïve mice (unpaired t-test, $p < 0.0001$).

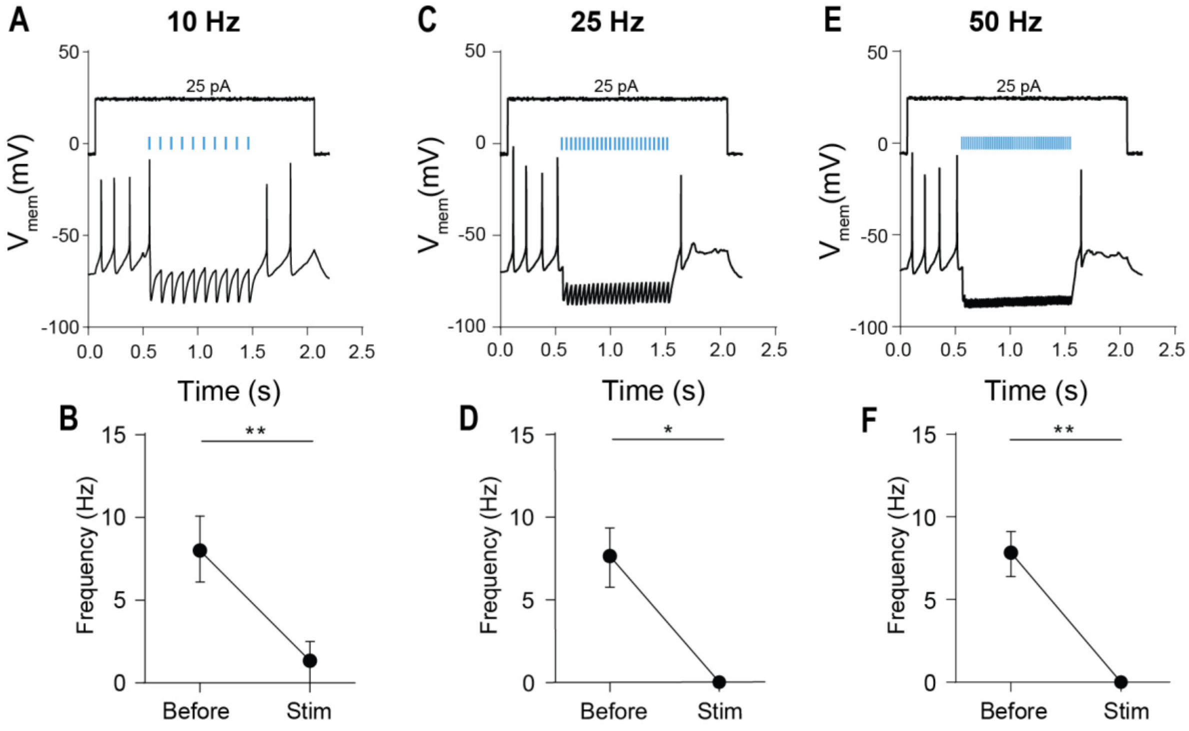

**Fig. S6.** ***In Vitro*** **current clamp recording confirming StGtaCR2 inhibition**
**A**) Example neuron showing inhibition of VTA DA neurons using 10 Hz stimulation with 473 nm light. Cellular membrane current was clamped with an injection of 25 pA of electrical current. **B**) Stimulation at 10 Hz significantly reduced firing frequency (paired t-test, $p < 0.01$). **C**) Example neuron showing inhibition of VTA DA neurons using 25 Hz stimulation with 473 nm light. **D**) Stimulation at 25 Hz significantly reduced firing frequency (paired t-test, $p < 0.05$). **E**) Example neuron showing inhibition of VTA DA neurons using 50 Hz stimulation with 473 nm light. **F**) Stimulation at 50 Hz significantly reduced firing frequency (paired t-test, $p < 0.01$). * $p < 0.05$; ** $p < 0.01$. Error bar indicates SEM.

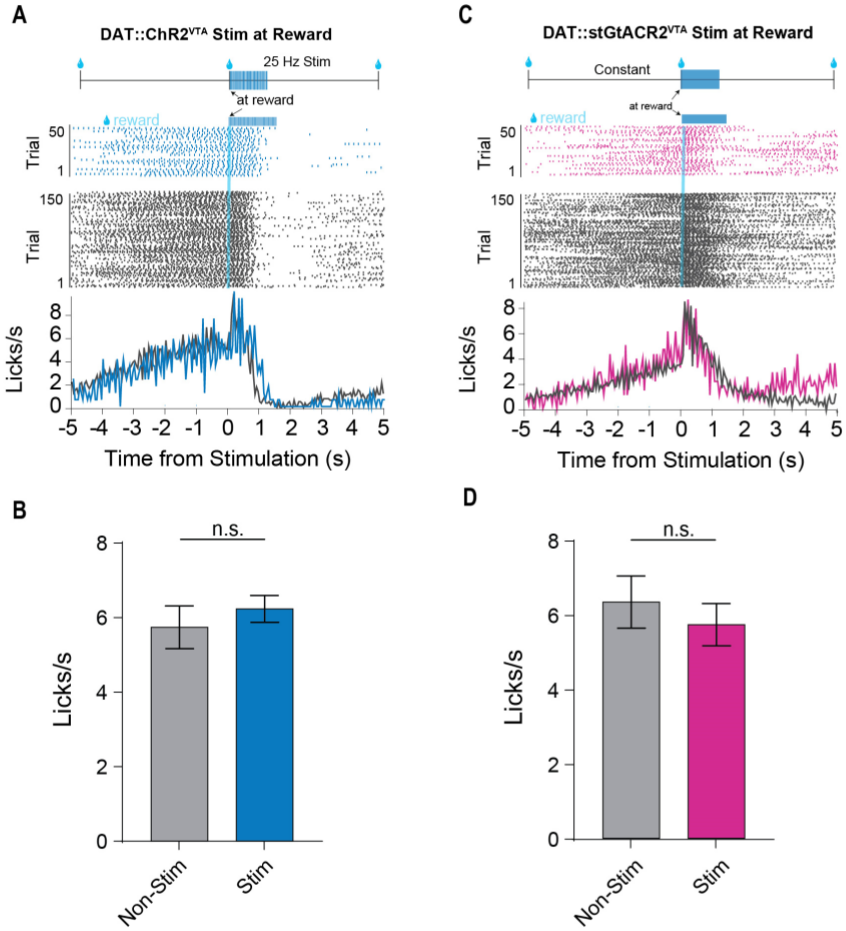

**Fig. S7. Optogenetic excitation or inhibition of VTA DA neurons has no effect on consummatory licking**

**A**) *Top:* Schematic of excitation of VTA DA neurons (DAT::ChR2VTA) at the time of reward delivery. *Bottom:* Optogenetic stimulation does not affect consummatory licking. Peri-event raster plots of licking during non-stimulation trials (grey) and stimulation trials (blue) for representative animal. Each line represents one trial. Below raster plots are average traces of lick rate for the data shown in the rasters. **B**) Optogenetic excitation VTA DA neurons during reward consumption has no effect on licking (paired t-test, $p > 0.05$). *Top:* Schematic of VTA DA inhibition at reward time *Bottom:* Optogenetic inhibition does not affect consummatory licking. Peri-event raster plots of licking during non-stimulation trials (grey) and stimulation trials (blue) for representative animal. **D**) Optogenetic inhibition of VTA DA neurons during reward consumption has no effect on licking (paired t-test, $p > 0.05$). Error bars indicate SEM.

**Movie S1. Stimulation of VTA DA neurons results in the generation of force in the forward direction along with licking in a reward-timing task**

Video shows a representative trial in which DAT::ChR2$_{VTA}$ stimulation results in forward force generation and anticipatory licking. Video is first played at normal speed then repeated in 0.25x time. Laser stimulation was 40 pulses, delivered 6 seconds prior to reward delivery at 20 Hz, and is indicated with the blue square.

**Movie S2. Single light pulse stimulation of VTA DA neurons is sufficient to produce forward force**

Video shows forward force generation and anticipatory licking following single light pulse stimulation of VTA DA neurons in DAT::ChR2$_{VTA}$ mice. Stimulation occurred during the fixed-time task at 6 seconds prior to reward. The same video is shown twice, first at normal speed, then repeated in 0.25x time. The blue rectangle presented in the slow-motion replay indicates time of single-light pulse delivery.
